# Energy deficit-independent stress response in the Frataxin-depleted heart: evidence that Integrated Stress Response can predominate over mTORC1 activation

**DOI:** 10.1101/2020.06.12.148361

**Authors:** César Vásquez-Trincado, Monika Patel, Aishwarya Sivaramakrishnan, Carmen Bekeová, Lauren Anderson-Pullinger, Nadan Wang, Hsin-Yao Tang, Erin L. Seifert

## Abstract

Friedreich’s ataxia (FRDA) is an inherited disorder caused by depletion of frataxin (FXN), a mitochondrial protein required for iron-sulfur cluster (ISC) biogenesis. Cardiac dysfunction is the main cause of death. Yet pathogenesis, and, more generally, how the heart adapts to FXN loss, remain poorly understood, though are expected to be linked to an energy deficit. We modified a transgenic (TG) mouse model of inducible FXN depletion that permits phenotypic evaluation of the heart at FXN levels < 20% of normal, without heart failure. We investigated substrate-specific bioenergetics and nutrient and stress signaling in the heart, to evaluate how this model responds to FXN depletion. After > 8 weeks with FXN levels < 20% of normal, TG hearts did not display overt hypertrophy and were in fact smaller; global protein translation was lower, while protein degradative pathways were unaltered. Cardiac contractility was maintained, likely due to preserved β-oxidation, though oxidative phosphorylation capacity for pyruvate was lower. Bioenergetics alterations matched mitochondrial proteomics changes, including a non-uniform decrease in abundance of ISC-containing proteins. In parallel, both mTORC1 signaling and the integrated stress response (ISR) were activated. The lack of overt cardiac hypertrophy, consistent with lower global protein translation, suggests that ISR predominated over mTORC1 activation. Suppression of a major ATP demanding process could benefit the FXN-depleted heart, at least short term. Thus, the FXN-depleted heart may enter a protective state, not necessarily linked to a major energy deficit. Finally, we propose the model used here as a pre-clinical model of cardiomyopathy in FRDA.

## INTRODUCTION

Friedreich’s Ataxia (FRDA) is caused by an expansion repeat of GAA between exon 1 and 2 of the gene encoding Frataxin (FXN). This leads to depletion of FXN which is part of the protein machinery responsible for iron-sulfur (Fe-S) cluster (ISC) biogenesis located in the mitochondrial matrix. It is generally accepted that FXN depletion below ∼30% of normal levels leads to pathology, the severity of which is generally correlated with GAA expansion length (1). FRDA is characterized by a fully penetrant ataxia, as well as other pathology, such as cardiomyopathy and progression to heart failure, that are less well predicted by GAA repeat length (2–5); thus cardiomyopathy in FRDA has been described as highly variable (5).

Cardiac dysfunction is the main cause of mortality in FRDA (3, 5). Yet, its pathogenesis, including a variable presentation, is not well understood, though it might be expected that a major energy deficit, due to loss of oxidative phosphorylation (oxphos) capacity, is a driver of pathology. Energy deficit seems logical as a source of pathology since the heart has an incessant requirement for ATP supplied by oxphos, and three of the electron transport chain (ETC) complexes require Fe-S centers for electron transfer, as do other enzymes of substrate metabolism within the mitochondrial matrix. Yet substrate oxidation pathways have not, to our knowledge, been studied in detail in cellular or animal models of FRDA.

An elegant time course study using the MCK (creatine kinase, muscle isoform) mouse model of FXN loss points to the activation of the integrated stress response (ISR) as an event concurrent with or preceding a putative energy deficit in the heart (6). The ISR involves phosphorylation of eIF2α (p-eIF2α) that leads to decreased global translation on one hand, and, on the other hand, to the translation of the transcription factor ATF4 and genes induced by ATF4 (7, 8). Induction of ATF4 targets can serve as readout of elevated ATF4 and is observed in many models of mitochondrial diseases (9–15). Considering models of mitochondrial disease, a small number of cell models have report a rise in p-eIF2α (13, 16), whereas most mouse models that investigated changes in signaling in response to mitochondrial dysfunction have focused on mTORC1 (17) (12, 18, 19), and AMPK signaling (14, 20). AMPK and mTORC1 signaling have received less attention in FRDA models, including the MCK mouse. Of note, the MCK model features cardiac hypertrophy (6, 21–23), which is not easily explained by elevated p-eIF2α. The latter point, together with the mitochondrial disease literature, suggests that multiple signaling pathways can be altered to drive phenotypes. Thus, for any given model, it would be useful to more broadly understand nutrient and stress signaling. A broader understanding might also expand the therapeutic options and provide insight into variability in the cardiac status of FRDA patients.

The MCK mouse model has been used to evaluate potential therapies (6, 21, 24) (25, 26). The model features a complete loss of FXN soon after birth, and rapid progression to heart failure. Yet, given the variability of cardiomyopathy in FRDA, it can be valuable to have available and to characterize other mouse models in which FXN protein is depleted by > 70%. Such a model was recently developed, with FXN depletion induced by Doxycycline (Doxy) (27). The mice developed a cardiomyopathy that was milder and slower in onset compared to the MCK mouse. Here we have used this newer genetic model to undertake a detailed study of substrate-specific bioenergetics in heart mitochondria and to broadly interrogate nutrient and stress signaling pathways. We used a modified Doxy dosing regimen that lowered mortality, and allowed phenotypes to be evaluated with FXN protein just below 20% of normal levels, and when FXN protein was fully depleted but before mice had undergone a major loss in body weight. We show that ∼20% FXN is sufficient to maintain heart function and oxphos capacity, with no evidence of altered signaling. When FXN was fully depleted, heart function and bioenergetics from β-oxidation were preserved, but hearts were smaller, AMPK and mTORC1 signaling were altered, and ISR activation was associated with lower global translation. We interpret this response as protective for the heart, at least in the short term, and propose this model as a pre-clinical model of FRDA.

## RESULTS

### Model of adult-onset doxycycline-induced FXN depletion

We have used a genetic mouse model to deplete FXN via doxycycline (Doxy) induction of the expression of an shRNA targeting *Fxn* mRNA. This is the same genetic model as that used by Chandran and colleagues (27). As done previously, Doxy dosing was initiated when mice were ∼9 weeks (wks) of age. However, we used a different dosing regimen; we supplied Doxy in the chow (200 p.p.m.) (Fig. 1A), whereas, in the earlier study, Doxy was supplied by intraperitoneal injection (5 mg/kg, with a 3-wk span at 10 mg/kg) and drinking water (2 mg/ml, daily). Both regimens resulted in complete FXN loss by 18 wks of Doxy treatment (see Fig. 1B).

**Figure 1:**
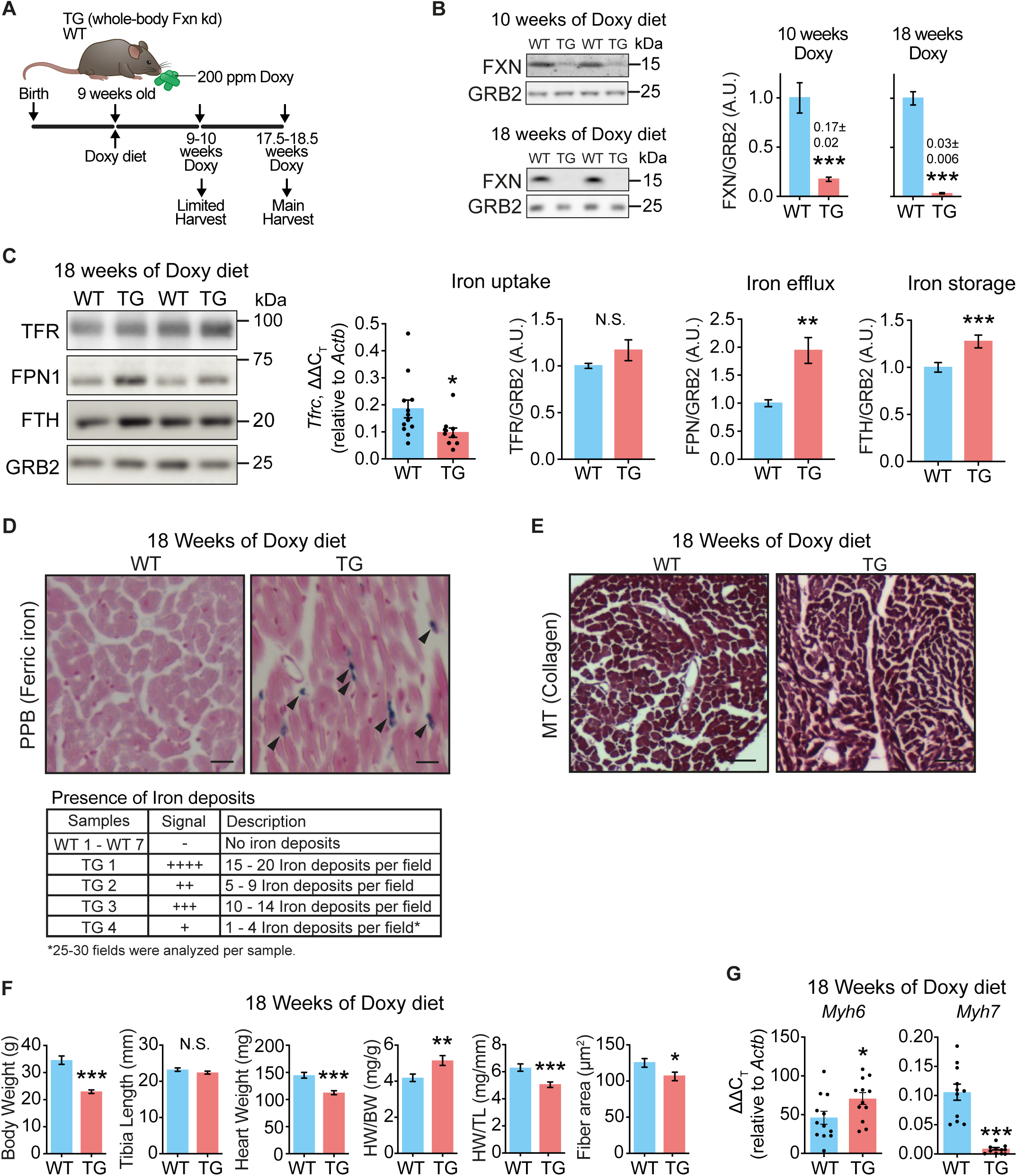
FXN-depleted hearts exhibit normal contractility but are smaller. **(A)** Experimental model. Mice harbored a whole body-wide doxycycline-inducible shRNA construct targeting *Fxn* (TG mice). Doxycycline (Doxy, 200 p.p.m.) was delivered in the chow. Both TG and wild-type mice (WT litter mates) received the Doxy-dosed diet. **(B)** Representative immunoblots of FXN and GRB2 (loading control) in heart lysates from WT and TG mice, and quantification (n=7/genotype). **(C)** Iron homeostasis in heart lysates, determined by immunoblot (left panels: TFR, FPN1, FTH, with GRB2 as loading control), and qPCR (*Tfrc*; ΔΔCT, relative to *β-actin*). Bar charts: mRNA levels of *Tfrc* (n=12 WT/10 TG, \*\**p*<0.01), and protein levels of TFR, FPN1 and FTH (n=4-7 WT/5 TG). **(D)** Cross-sections from heart stained with Perls’ Prussian Blue (PPB) to reveal iron deposition. Table: Qualitative score of PPB-stained sections to evaluate abundance of iron deposits (n=7 WT/ 4 TG). **(E)** Cross-sections from hearts stained with Masson’s trichrome to evaluate fibrosis. Fibrosis was not detected (n= 7 WT/4 TG). **(F)** Body weight, tibia length (TL), heart weight (HW), tibia length to heart weight ratio (HW/TL) (n=7/genotype), and cross-sectional area of heart muscle fibers (n= 7 WT/4 TG). **(G)** Transcript levels (ΔΔCT, relative to *β-actin*) of *Myh6* (α-MHC) and *Myh7* (β-MHC) (n=11-12/genotype; points are values from each sample). Panels B, C, E-G: values are mean ± s.e.m. Statistical comparison was by unpaired t-test; *p<0.05, **p<0.01, ***p<0.001. N.S.: not significant.

Wild-type (WT) mice and mice harboring one copy of the Doxy-inducible shRNA transgene against *Fxn* (TG) were generated from a WT X TG cross (27). TG and WT littermates were fed the Doxy diet. Gross phenotypic changes were evident in TG mice after ∼13 wks of Doxy feeding. Notably, whereas body weight of TG and WT mice was comparable during the first 13 wks of Doxy, TG mice subsequently underwent a progressive weight loss that was pronounced by 20 wks of Doxy (SFig 1A), similar to that reported by Chandran (27). Whereas mortality in TG studied by Chandran was evident by 12 wks of Doxy (27), some mortality was evident only after ∼16 wks of Doxy feeding, and by 18 wks of Doxy, ∼35% of the TG mice had died (SFig. 1B), similar to what Chandran observed at 12 wks of Doxy. The cause of death was not obvious, but was not correlated with body weight loss.

We focused our studies on two time points: Mice fed Doxy for 9-10 wks (“10 wks”) (substantial FXN protein loss in the heart, see Fig. 1B) and for 17.5-18.5 wks (“18 wks”) (full FXN loss in the heart (Fig. 1B), some body weight loss, but prior to major body weight loss and mortality).

Focusing on the heart, after 10 wks of Doxy feeding, FXN was ∼80% depleted in the heart of TG mice (Fig. 1B). By 18 wks of Doxy, FXN was no longer detectable in TG hearts, as assessed by immunoblot (Fig. 1B) or by mass spectrometry (see Supplemental Table). Mass spectrometry analysis of isolated heart mitochondria from 18 wk Doxy-fed mice failed to identify unique FXN peptides in heart mitochondria samples from TG mice whereas multiple peptides were identified in samples from WT heart mitochondria (see Supplemental Table).

### Absence of overt pathology in FXN-depleted hearts, however heart mass was lower

Next we evaluated the levels of iron handling proteins in the heart as well as iron deposition, because marked changes in these proteins occur in FXN-depleted mouse hearts, and iron deposits have been reported in FXN-depleted mouse and human hearts (reviewed in (28)). Hearts from TG mice fed Doxy for 18 wks showed lower levels of Transferrin Receptor mRNA and unchanged protein levels, accompanied by higher levels of iron efflux and storage proteins, Ferroportin and Ferritin respectively (Fig. 1C). To estimate iron abundance in the heart, Perls’ staining was used which measures ferric iron, and includes iron bound by Ferritin (but not heme iron); Perls’ staining revealed iron deposits in TG but not WT hearts from 18-wk Doxy-fed mice (Fig. 1D). Thus TG hearts responded to FXN depletion with an iron homeostatic response consistent with elevated cellular iron (29).

The elevated ferric iron in TG hearts is abnormal and could be accompanied by pathological morphological and functional changes in the heart. Yet, histological evaluation of WT and TG hearts from 18-wk Doxy-fed mice revealed normal cardiac fiber structure and absence of fibrosis (Fig. 1E). Furthermore, echocardiography to evaluate cardiac function revealed slightly higher fractional shortening in hearts from TG mice fed Doxy for 18 wks, a change already evident in 10-wk Doxy-fed TG mice (Table 1). The greater fractional shortening was driven primarily by a smaller left ventricular end-systolic diameter (Table 1), consistent with increased contractility.

**Table 1.**
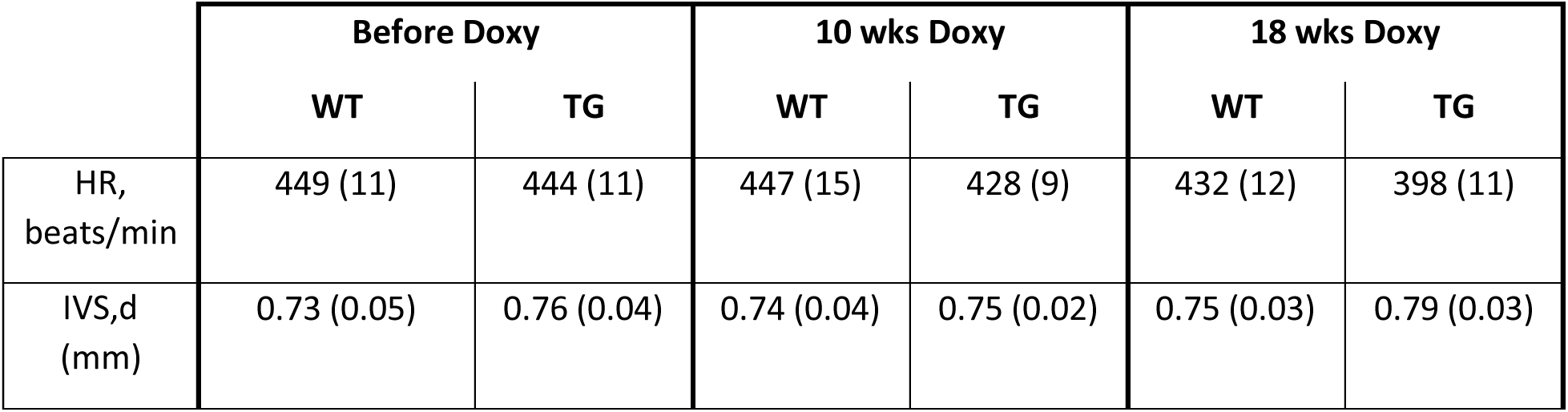
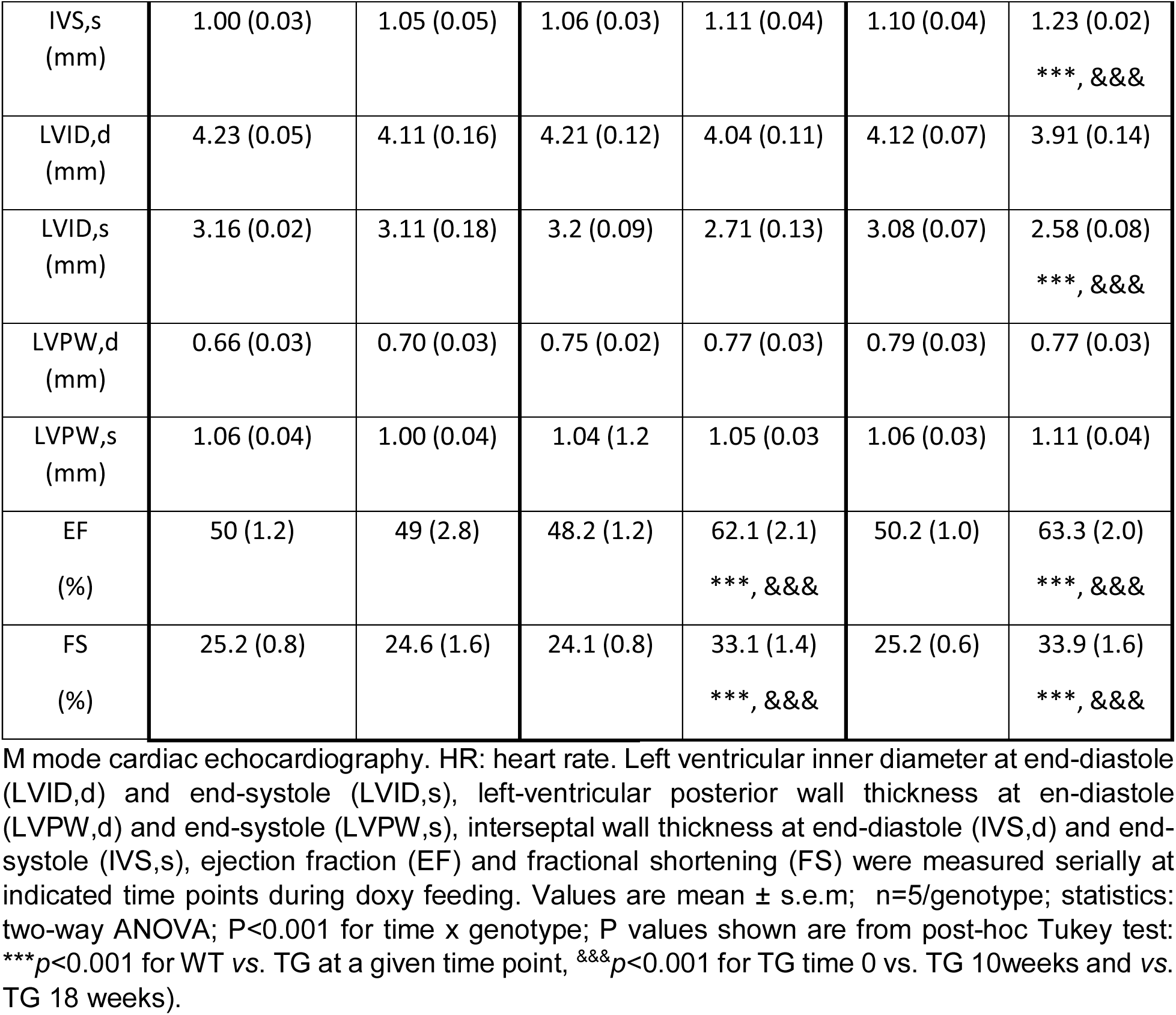
Echocardiography.

|  | Before Doxy |  | 10 wks Doxy |  | 18 wks Doxy |  |
| --- | --- | --- | --- | --- | --- | --- |
|  | WT | TG | WT | TG | WT | TG |
| HR, beats/min | 449 (11) | 444 (11) | 447 (15) | 428 (9) | 432 (12) | 398 (11) |
| IVS,d (mm) | 0.73 (0.05) | 0.76 (0.04) | 0.74 (0.04) | 0.75 (0.02) | 0.75 (0.03) | 0.79 (0.03) |
| IVS,s<br>(mm) | 1.00 (0.03) | 1.05 (0.05) | 1.06 (0.03) | 1.11 (0.04) | 1.10 (0.04) | 1.23 (0.02)<br>***, &&& |
| LVID,d<br>(mm) | 4.23 (0.05) | 4.11 (0.16) | 4.21 (0.12) | 4.04 (0.11) | 4.12 (0.07) | 3.91 (0.14) |
| LVID,s<br>(mm) | 3.16 (0.02) | 3.11 (0.18) | 3.2 (0.09) | 2.71 (0.13) | 3.08 (0.07) | 2.58 (0.08)<br>***, &&& |
| LVPW,d<br>(mm) | 0.66 (0.03) | 0.70 (0.03) | 0.75 (0.02) | 0.77 (0.03) | 0.79 (0.03) | 0.77 (0.03) |
| LVPW,s<br>(mm) | 1.06 (0.04) | 1.00 (0.04) | 1.04 (1.2) | 1.05 (0.03) | 1.06 (0.03) | 1.11 (0.04) |
| EF<br>(%) | 50 (1.2) | 49 (2.8) | 48.2 (1.2) | 62.1 (2.1)<br>***, &&& | 50.2 (1.0) | 63.3 (2.0)<br>***, &&& |
| FS<br>(%) | 25.2 (0.8) | 24.6 (1.6) | 24.1 (0.8) | 33.1 (1.4)<br>***, &&& | 25.2 (0.6) | 33.9 (1.6)<br>***, &&& |

M mode cardiac echocardiography. HR: heart rate. Left ventricular inner diameter at end-diastole (LVID,d) and end-systole (LVID,s), left-ventricular posterior wall thickness at end-diastole (LVPW,d) and end-systole (LVPW,s), interseptal wall thickness at end-diastole (IVS,d) and end-systole (IVS,s), ejection fraction (EF) and fractional shortening (FS) were measured serially at indicated time points during doxy feeding. Values are mean ± s.e.m; n=5/genotype; statistics: two-way ANOVA; P<0.001 for time x genotype; P values shown are from post-hoc Tukey test: \*\*\*p<0.001 for WT vs. TG at a given time point, &&&p<0.001 for TG time 0 vs. TG 10weeks and vs. TG 18 weeks).

Cardiac hypertrophy is often a feature of the cardiomyopathy of FRDA (Tsou et al., 2011). Thus we determined if cardiac hypertrophy was present in TG hearts. Morphometric evaluation of the heart histology revealed a smaller cross-sectional area of fibers in TG hearts from 18-wk Doxy-fed mice (Fig. 1F). Furthermore, measurement of heart weight in 18-wk Doxy-fed mice revealed a lower heart weight/tibia length ratio (HW/TL) in TG mice, though HW relative to body weight was higher (Fig. 1F). We also noted that LV posterior wall and septal wall thickness were unchanged in TG hearts (Table 1). To determine if smaller heart weight is intrinsic to the TG genotype, we measured HW/TL in mice that were not fed the Doxy diet and found no difference between WT and TG (SFig. 1C). These data show that FXN-depleted TG hearts are not overtly hypertrophic, and, in fact, are smaller than the heart of Doxy-fed WT mice. The data also suggest some preservation of LV mass in FXN-depleted TG hearts.

Lower levels of *Myh6* (α-MHC; myosin heavy chain, α isoform) and higher levels of *Myh7* (β-MHC) are often found in diseased hearts (30). However, TG hearts from 18-wk Doxy-fed mice showed slightly elevated *Myh6* and lower *Myh7* (Fig. 1G).

The expression pattern of the MHC isoforms, together with the functional and morphological data, indicated a lack of overt pathology in FXN-depleted TG hearts, even after 18 wks of FXN depletion. However, the data also suggest that FXN-depleted hearts were not normal. We hypothesized that the status of TG hearts reflected adaptive changes. To address this hypothesis, we questioned how, energetically, the FXN-depleted hearts could sustain their function (Fig. 2, Fig. 3), and whether the sustained function and smaller heart size might reflect an altered pattern of stress and nutrient signaling (Fig. 4-6).

**Figure 2:**
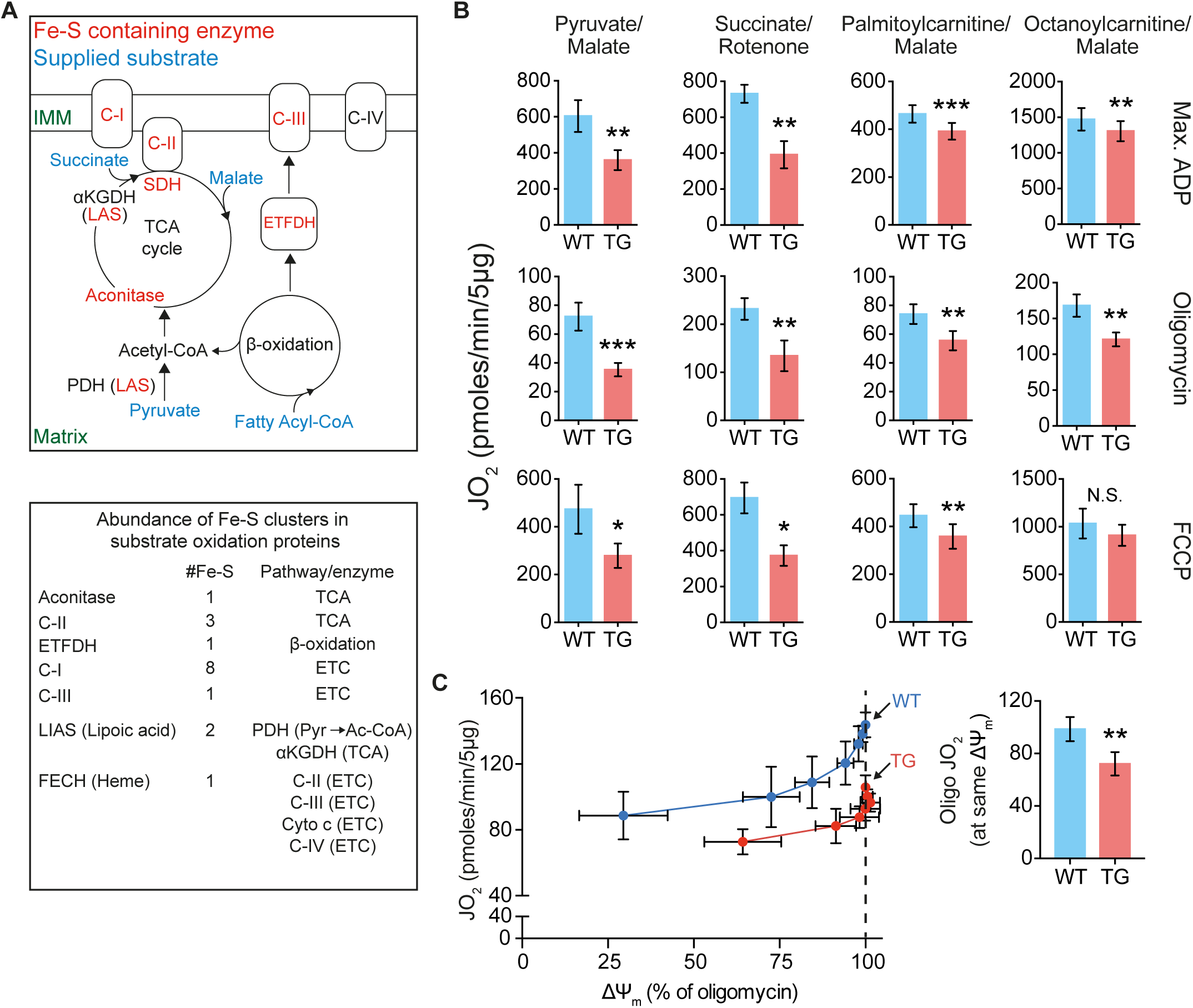
Substrate-dependent decrease in mitochondrial oxphos in TG hearts. All measurements were done in heart mitochondria from WT and TG mice fed Doxy for 18 wks. **(A)** *Upper*: Diagram shows Fe-S cluster-containing enzymes (red), and the substrates supplied in bioenergetics experiments (blue). *Lower*: List of Fe-S cluster-containing complexes/enzymes with corresponding number of Fe-S clusters and associated metabolic pathway or enzyme. ETFDH: Electron transfer flavoprotein dehydrogenase, LIAS: Lipoic acid synthetase, FECH: Ferrochelatase, PDH: Pyruvate dehydrogenase, αKGDH: α-ketoglutarate dehydrogenase, cyto c: Cytochrome c. C-I, C-II, CIII and C-IV are mitochondrial respiratory complexes I, II, III and IV. Pyr: Pyruvate; Ac-CoA: Acetyl-CoA. **(B)** Oxygen consumption rate (JO2) measured in isolated heart mitochondria supplied with pyruvate/malate (10 mM/5 mM), succinate (10 mM + 1 µM rotenone to prevent electron backflow through complex I), palmitoyl-L-carnitine plus malate (20 µM + 1 mM), or octanoyl-L-carnitine plus malate (200 µM+1 mM). Max. ADP, saturating [ADP] (4 mM), was used to evaluate JO2 reflecting maximal oxphos. Oligomycin was used to evaluated JO2 that reflects maximal leak-dependent oxidation. The chemical uncoupler, FCCP (1 µM), 2.5 µg/ml) as used to evaluate JO2 that reflects maximal electron transport chain capacity under the prevailing substrate conditions. N=6/genotype except for octanoyl-L-carnitine (n=4/genoype). **(C)** Proton leak was determined by measuring mitochondrial membrane potential (ΔΨm) and JO2 during antimycin (Complex III inhibitor) titration of octanoyl-L-carnitine oxidation, in the presence of oligomycin. *Bar chart:* JO2 values at the same ΔΨm (dashed line in the panel at left) (n=3/genotype). Panels B and C: values are mean ± s.e.m. Statistical analysis was by unpaired t-test; *p<0.05, **p<0.01. N.S.: not significant.

**Figure 3:**
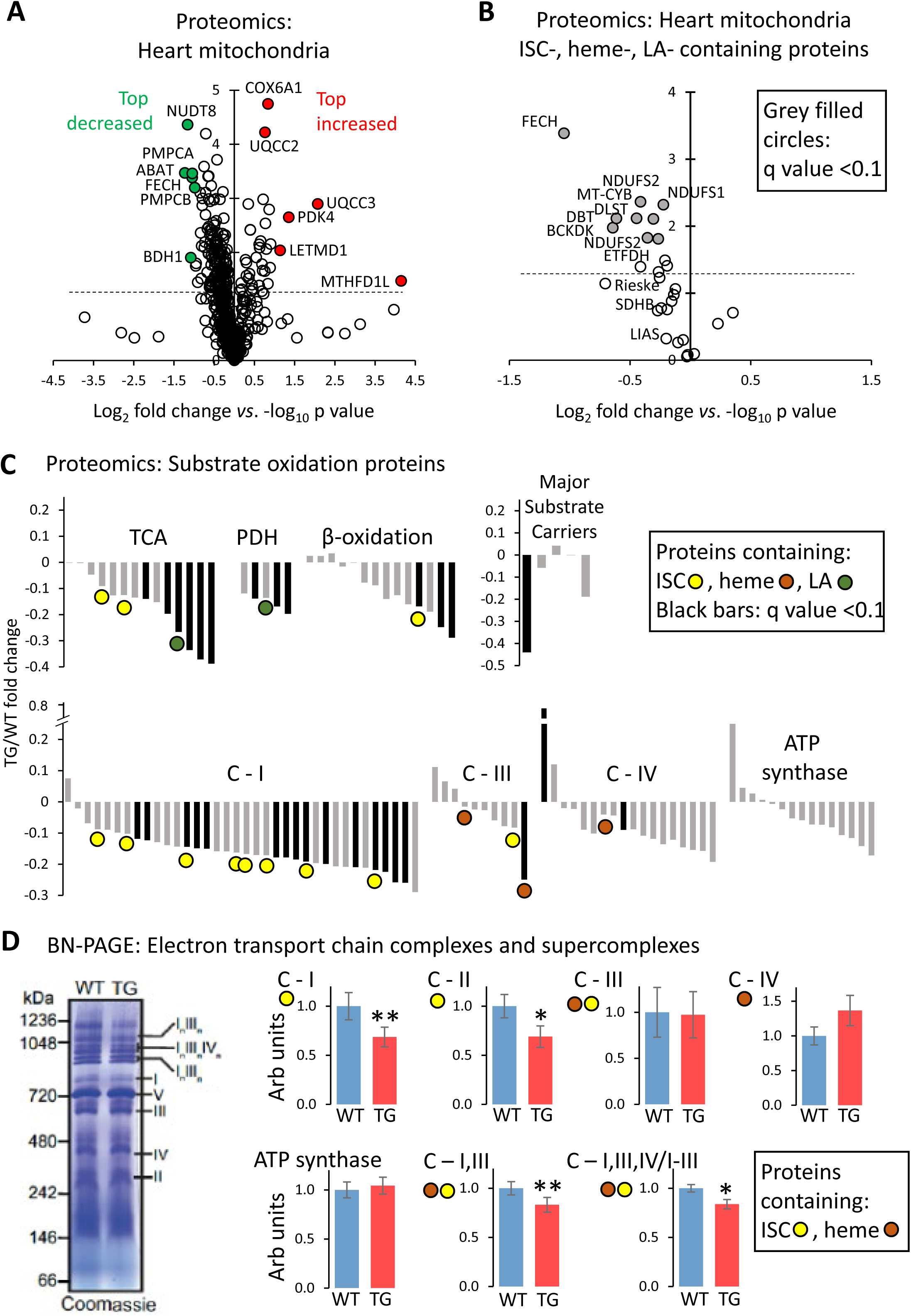
Proteomics analysis of heart mitochondria: variable effect of FXN depletion on ISC-containing proteins. All measurements were done in heart mitochondria from WT and TG mice fed Doxy diet for 18 wks. **(A)** Volcano plot (log2 TG/WT fold change *vs*. log10 p value, unpaired t-test) of heart mitochondrial proteins determined by mass spectroscopy (n=3/genotype). Inclusion criteria: proteins identification based on > 1 peptide in at least 1 genotype. Dashed line: p=0.05 (unpaired t-test). **(B)** Volcano plot (log2 TG/WT fold change *vs*. log10 p value, unpaired t-test) of heart mitochondrial proteins containing iron-sulfur clusters (ISC), or heme or lipoic acid (LA) moieties. Proteins determined by mass spectroscopy (n=3/genotype). Dashed line: p=0.05 (unpaired t-test). Black-filled circles: proteins with q value (p value corrected for false discovery rate) < 0.10. **(C)** Fold change of substrate oxidation-related proteins, determined by mass spectroscopy (n=3/genotype). Black bars: proteins with q value < 0.10. TCA: tricarboxylic acid cycle; PDH: pyruvate dehydrogenase complex; C-I,III,IV: complex I, III, IV of the electron transport chain. Complex II subunits are included in the “TCA” category. **(D)** BN-PAGE gel electrophoresis of isolated heart mitochondria, to reveal the abundance of electron transport chain (ETC) complexes and supercomplexes. *Left*: Example of Coomassie-stained gel, with ETC complexes and supercomplexes shown by Roman numerals. *Right*: Quantification; values are mean ± s.e.m (n=5/genotype, *p<0.05, **p<0.01 by unpaired t-test).

**Figure 4:**
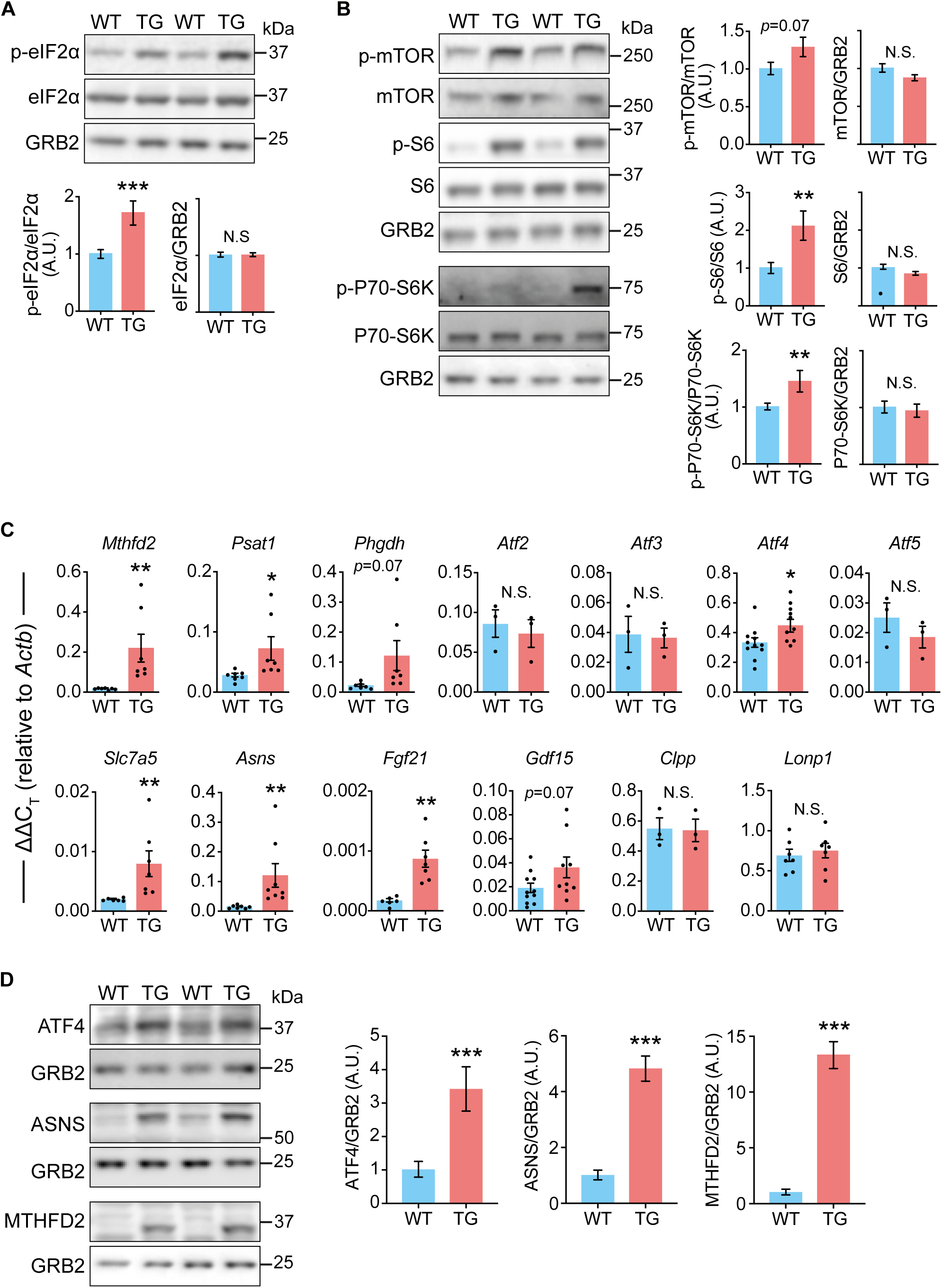
Elevated mTORC1 signaling and integrated stress response activation in the heart after 18 weeks of FXN depletion. All measurements were done in heart lysates from WT and TG mice fed Doxy for 18 wks. **(A)** Phosphorylation status of eIF2α. *Top*: Representative immunoblot of p-eIF2α(Ser52), total eIF2α and GRB2 from heart lysates of WT and TG mice. Lower: Quantification (n=9 WT/7 TG). **(B)** mTORC1 signaling. *Left*: Representative immunoblots of p-mTORC1 (Ser2448), total mTOR, p-S6 (Ser235/236), total S6, p-P70S6K, total P70S6K, and GRB2 and GAPDH (loading controls). *Right*: quantification (n=9 WT/7 TG). **(C)** Transcripts levels (ΔΔCT; relative to *Actb* (β-Actin)) of stress signaling targets. Points show values from each sample. **(D)** Protein levels of ATF4 and targets. Left: Representative immunoblots. GRB2 was used as loading control. *Right:* Quantification (n=6/genotype). In all panels: bars represent mean ± s.e.m. Statistical comparison was by unpaired t-test, \**p*<0.05, \*\**p*<0.01. N.S.: not significant.

**Figure 6:**
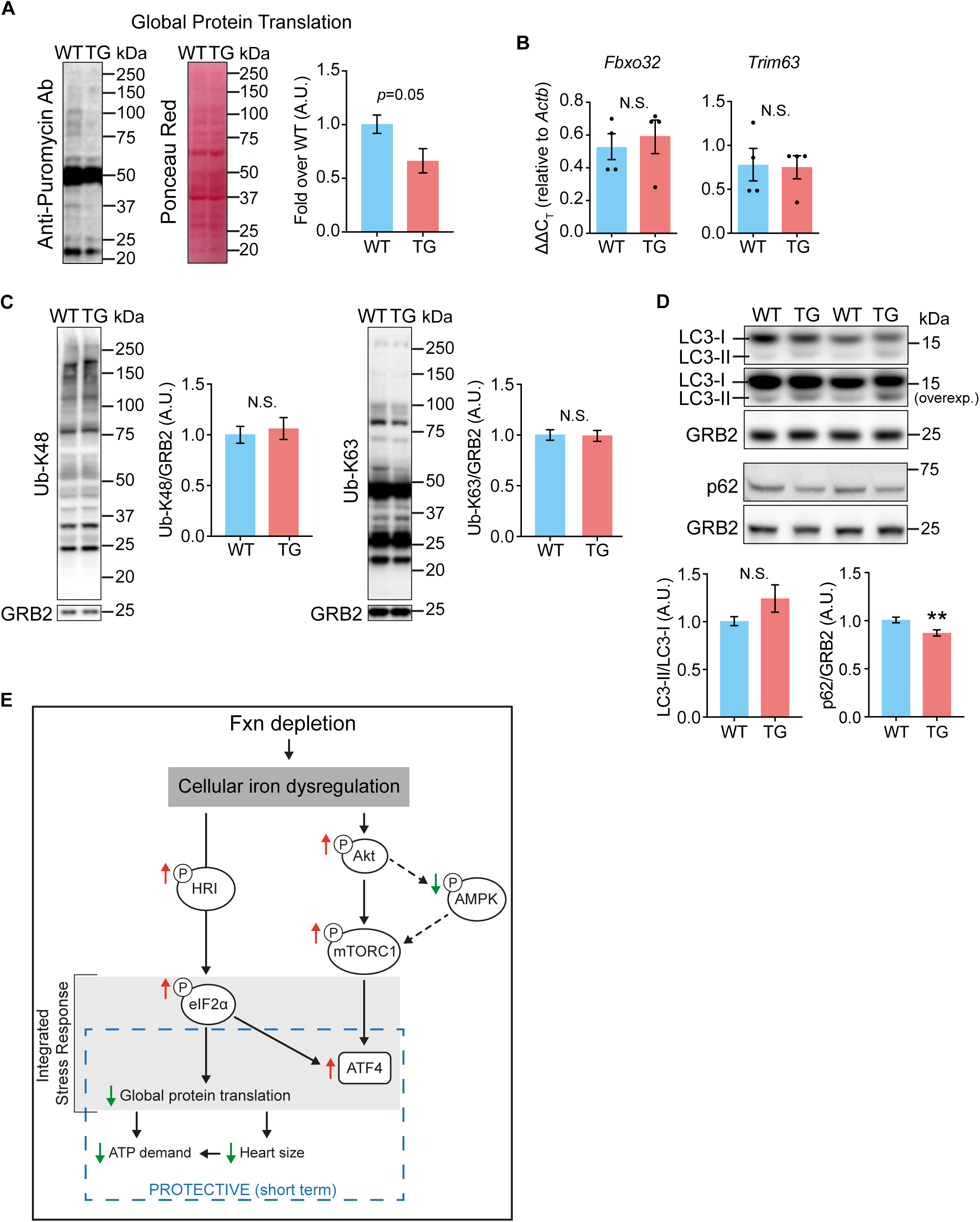
Decreased global protein translation in hearts depleted of FXN for 18 wks. **(A)** Global protein translation was determined using the SUnSET method. *Left*: Representative immunoblot using anti-puromycin antibody, with Ponceau Red staining to estimate protein loading. *Right*: Quantification of puromycin signal (*p<0.05 by unpaired t-test, n=4/genotype). **(B), (C)** Protein degradation, estimated by transcript levels of *Fbxo32* and *Trim63* (Panel B, n=4/genotype, not significant (N.S.), by unpaired t-test), and by K48-linked ubiquitination of proteins by immunoblot (Panel C; N.S.: not significant by unpaired t-test, n=6/genotype). **(D)** Autophagy, estimated by levels of LC3 and lipidated LC3, and levels of p62, all by immunoblot. N.S.: not significant by unpaired t-test, n=4/genotype. **(E)** Model of altered signaling in FXN-depleted hearts.

### The extent of oxphos suppression in TG heart mitochondria is substrate-dependent

Many of the Fe-S cluster (ISC)-containing proteins within the mitochondrial matrix participate in substrate oxidation that is necessary for oxidative ATP synthesis (Fig. 2A), an absolute requirement for heart function. In particular, ISC-containing proteins are present in the tricarboxylic acid (TCA) cycle (specifically, aconitase and succinate dehydrogenase (SDH)), Complexes I and III of the electron transport chain (ETC), and in the electron transfer flavoprotein dehydrogenase (ETFDH) that directly transfers electrons from the acyl-CoA dehydrogenase-coupled FAD^+^ reduction of β-oxidation to Complex III via ubiquinol. Lipoic acid synthetase (LIAS) in the mitochondrial matrix also contains an ISC and is needed to synthesize the lipoic acid moiety found in the E2 subunit of PDH and α-KGDH (pyruvate dehydrogenase and α-ketoglutarate dehydrogenase, respectively). Finally, matrix-localized ferrochelatase (FECH) is the main rate-controlling enzyme in heme biosynthesis; among the substrate oxidizing proteins, heme is found in subunits of Complex II, III and IV, and in cytochrome c. ISCs are thus required for the oxidation of fatty acids and pyruvate, the substrates normally utilized by the heart to generate ATP.

Thus, FXN depletion was expected to be accompanied by a decrease in oxidative phosphorylation (oxphos) capacity in heart mitochondria, whether supplied with pyruvate or fatty acids. To this end, we performed a detailed bioenergetics analysis in isolated heart mitochondria supplied with four substrates: pyruvate (with malate), succinate (with rotenone to inhibit backflow of electrons to Complex I) and two fatty acid substrates (palmitoyl-L-carnitine (PCarn) and octanoyl-L-carnitine (OCarn), each with malate). To oxidize these substrates, several ISC-containing proteins are needed, and some of these are substrate-specific (Fig. 2A). Moreover, oxidation of pyruvate relies on more ISC-containing proteins than does fatty acid-derived electron supply to Complex III via the ETFDH and ubiquinol (Fig. 2A). Oxidation was evaluated as O2 consumption rate (JO2) under the following conditions: saturating concentrations of substrate and ADP were used to evaluate maximal oxphos; oligomycin, an ATP synthase inhibitor, was used to determine maximal leak-dependent JO2; and the chemical uncoupler, FCCP, was used to evaluate maximal ETC capacity.

In heart mitochondria from TG mice fed Doxy for 10 wks (∼80% FXN loss), with pyruvate/malate as the substrate, ADP-stimulated oxphos tended to be lower and leak-dependent JO2 was significantly lower (SFig. 2). However, for succinate or PCarn/malate, maximal oxphos, leak and ETC capacity were similar between WT and TG mitochondria (SFig. 2). After 18 wks of Doxy (complete FXN loss in TG hearts) TG mitochondria supplied with pyruvate/malate or succinate showed a pronounced (40-50%) deficit in maximal oxphos, leak-dependent JO2 and maximal ETC capacity (Fig. 2B). In contrast, TG mitochondria supplied with PCarn/malate showed a modest (∼15%) decrease in these parameters (Fig. 2B).

We noted that maximal ADP-driven JO2 in WT mitochondria supplied with PCarn/malate was similar to the rate in TG mitochondria supplied with pyruvate/malate. Thus we questioned if the modest changes in bioenergetics parameters in TG mitochondria oxidizing PCarn/malate reflected a limitation of the PCarn/malate reaction. To test this possibility we used octanoyl-L-carnitine (OCarn) which, despite its shorter carbon chain length (C8:0), had higher maximal oxphos than PCarn (C16:0) likely because more substrate can be oxidized without causing uncoupling (31); indeed, the ratio of maximal oxphos to maximal leak JO2 was ∼5.5 for PCarn/malate and ∼8.8 for OCarn/malate in WT mitochondria. Yet, even with OCarn/malate, there was only a modest decrease in JO2 in TG mitochondria (Fig. 2B). In fact, though maximal oxphos of TG mitochondria oxidizing OCarn/malate was slightly (and significantly) less than in WT mitochondria, maximal oxphos of TG mitochondria oxidizing OCarn/malate was higher than when pyruvate/malate was supplied to TG mitochondria.

We noted that maximal leak-dependent (oligomycin-insensitive) JO2 was lower in TG heart mitochondria supplied with any of the substrates tested (see Fig. 2B). Leak-dependent JO2 can reflect supply of reducing equivalents to the ETC to generate the driving force for oxphos or can reflect proton leak back into the matrix by pathways other than the ATP synthase, lowering the efficiency of substrate oxidation to generate ATP. To discriminate between these possibilities, and thus to gain mechanistic insight, we evaluated leak-dependent JO2 as a function of driving force (membrane potential, ΔΨm), by titrating OCarn/malate oxidation with the Complex III inhibitor antimycin, in the presence of oligomycin. This experiment allows leak-dependent JO2 to be compared between TG and WT at the same driving force (i.e., supply of reducing equivalents). Consistently, JO2 was lower at any given ΔΨm in TG *vs*. WT heart mitochondria (Fig. 2C), indicating lower proton leak in TG mitochondria.

Altogether, the bioenergetics data indicate that, as expected, FXN loss leads to lower oxphos capacity. The decrease was evident when FXN was completely lost, and not when ∼20% of FXN remained (10 wks Doxy). The data also reveal a substrate-dependence of the lower oxphos capacity which, along with the lower proton leak in TG mitochondria, suggest that mitochondrial substrate metabolism in FXN-depleted mitochondria cannot simply be ascribed to changes in ISC-containing proteins.

### Broad decreases in levels of Complex I and TCA cycle enzymes in TG heart mitochondria

To address the differential impact of FXN loss on substrate oxidation, and to more broadly understand the impact of FXN depletion on heart mitochondria, we undertook a proteomics analysis of heart mitochondria isolated from 18-wk Doxy-fed WT and Tg mice. A total of 595 mitochondrial proteins were identified (Supplemental Table). A volcano plot of TG protein abundance relative to WT levels revealed that many proteins were less expressed in TG mitochondria (Fig. 3A, B); these included some, but not all, ISC-, lipoic acid, or heme-containing proteins, as well as other proteins. Among the most substantially decreased non-ISC-containing proteins were PMPCA and PMPCB, the α and β subunits of the matrix-localized Mitochondrial Processing Peptidase (MPP), a major protease that cleaves the N terminal mitochondrial targeting sequence of nuclear-encoded mitochondria proteins including FXN (32). Several proteins were increased in TG mitochondria; among the proteins showing the greatest increases was PDK4 (pyruvate dehydrogenase kinase) which phosphorylates and inhibits PDH (Fig. 3A).

Among the ISC-, lipoic acid- and heme-containing proteins, FECH, required for cellular heme synthesis, showed the greatest decrease, and was also one of the most substantially decreased protein we detected (Fig. 3A, B). Lower FECH expression was confirmed by immunoblot in heart lysates and isolated mitochondria (SFig. 3). Heme-containing matrix were however not uniformly decreased in abundance in TG mitochondria. Cytochrome b of Complex III was significantly decreased, whereas the cytochrome C1 subunits of Complex III and the mitochondrial-DNA-encoded mt-CO1 subunit of Complex IV were unchanged (Fig. 3C: brown dots). Cytochrome c was not detected by mass spectrometry, however immunoblotting revealed a robust decrease in expression in TG mitochondria (SFig. 3A). Besides FECH, other substantially decreased (q<0.1) ISC-containing proteins were subunits of Complex I (Fig. 3B). SDHB, aconitase and the Rieske protein of Complex III trended lower, whereas LIAS (required for lipoic acid synthesis) was clearly unchanged, as was ISC-containing NDUFS7 of Complex I (see Fig. 3C). Immunoblotting of heart lysates and isolated heart mitochondria confirmed lower SDHB whereas differences between WT and TG could not be detected in the abundance of aconitase, Rieske protein or LIAS (SFig. 3). Note that antibodies were first evaluated to determine conditions which that yielded a ∼linear dynamic range (not shown). Thus, ISC-containing proteins were not uniformly downregulated in mitochondria from FXN-depleted hearts. That the proteomics findings from isolated mitochondria could be confirmed in heart lysates indicates that the findings in isolated mitochondria do not reflect a sub-selection of the healthiest mitochondria.

We next focused on proteins of the main substrate oxidation pathways in the heart, particularly subunits of the ETC and ATP synthase, TCA cycle and β-oxidation proteins, subunits of PDH and substrate carriers. These are plotted in Fig. 3C, and include proteins containing ISCs (yellow dots), heme (brown dots) and lipoic acid (green). The most affected pathways or protein complexes were the TCA cycle (6/14 proteins decreased including SDHA, p < 0.05, q < 0.1), PDH (3/5 subunits decreased, p < 0.05, q < 0.1), and Complex I (15/36 subunits were decreased, p < 0.05, q < 0.1); immunoblotting of SDHA and NDUFA9 (Complex I subunit) also revealed significant decreases in these proteins in TG lysates and mitochondria (SFig. 3). Of the 24 members of these pathways determined by mass spectrometry to be decreased (q<0.1) in TG mitochondria, 4 contained ISCs (out of the 12 ISC proteins in these pathways), 1 contained heme (out of 3 heme-containing proteins), and 1 contained lipoic acid (of 2). Only one protein was increased (q<0.1; a subunit of Complex IV). Levels of transporters for the main substrates (MPC1/2 for pyruvate and CPT1a/b for activated long-chain fatty acids) were unchanged, though there was ∼50% lower abundance of the dicarboxylate carrier (SLC25A10) which transports malate and succinate into the matrix (Fig. 3C). The lower abundance of SLC25A10 did not reflect a broad decrease in SLC25A family members (see Supplemental Table).

In light of the decreased proton leak (Fig2C) and the recent report that proton leak is mediated by the adenine nucleotide translocase (ANT) (33), we noted the levels of ANT1 and ANT2, the two ANT isoforms that were detected. ANT1, the major isoform in the heart, showed similar expression in TG and WT mitochondria (p = 0.6). ANT2, detected at levels an order of magnitude lower than those of ANT1, was higher in TG mitochondria (p = 0.01; q = 0.1). Thus, ANT abundance likely do not explain the lower proton leak in TG mitochondria.

Beyond immunoblot detection in heart lysates to address possible sub-selection of healthy mitochondria, another quality-control issue we considered is whether there was biasing based on protein localization within mitochondria. Such biasing seems unlikely, since membrane-localized and peripheral-matrix localized subunits of Complexes I and III (34–36) were roughly equally likely to be found to be significantly changed in TG mitochondria.

Because ETC subunits function in complexes and higher order supercomplexes (SC), we performed blue native (BN)-PAGE analysis to determine complex SC abundance and activity in heart mitochondria from mice fed Doxy for 18 wks. The abundance of assembled Complex III and IV and ATP synthase was similar between WT and TG mitochondria (Fig. 3D). However, TG mitochondria exhibited slightly but significantly lower abundance of assembled Complex I and II (Fig3D) and in-gel activity of these complexes was also lower (SFig. 3D). The abundance of SCs, which contain Complexes I, III and IV, was also modestly lower in TG mitochondria (Fig. 3D).

To also test for a metabolic switch from β-oxidation to glycolysis, as described in hearts with compromised oxphos (37), we evaluated mRNA levels of transporters and enzymes involved in glucose transport and metabolism (*Slc2a4*, *Hk2*, *Pfkfb1*) and fatty acid transport and metabolism (*Cd36*, *Cpt1b*, *Acadm*) in hearts from 18-wk Doxy-fed mice. There were no differences between WT and TG hearts in the levels of any of these transcripts (SFig. 4).

Many mitochondrial proteins were expressed at lower abundance in hearts from 18-wk Doxy-fed TG mice. Yet, a few proteins were increased. In particular, MTHFD1L (methylene tetrahydrofolate dehydrogenase 1 like, a matrix protein involved in folate metabolism) was barely detectable in WT mitochondria (0 or 1 peptide was detected by mass spectrometry) but readily detected after FXN loss (4 or 5 peptides detected). MTHFD1L is a target of the transcription factor ATF4 (38). CYB5R1 was also substantially increased and is an ATF4 target (39). ATF4 expression is normally low but its translation increases when translation initiation factor eIF2α is phosphorylated which otherwise leads to global translation inhibition (8). ATF4 induces genes that counter cell stress. Decreased global protein translation also counters cell stress by lowering the demand for nutrients and energy. This response is referred to as the Integrated Stress Response (ISR) (8). ATF4 levels can also be induced via the mTORC1 pathway (40, 41). Thus, we evaluated the status of the ISC and mTORC1 signaling.

### Stress signaling and aberrant nutrient signaling in FXN-depleted hearts

We evaluated the phosphorylation of eIF2α, the core event of ISR activation (8), in heart lysates. Compared to WT hearts, TG hearts from 18-wk Doxy-fed mice showed substantially higher levels of phosphorylated eIF2α, with no changes in total eIF2α (Fig. 4A). Investigating the mTORC1 pathway, there was evidence for increased activity in TG hearts; though phosphorylation of mTOR showed no difference between WT and TG hearts (Fig. 4B), downstream components of this pathway, namely p-S6 and p-P70-S6K were more phosphorylated in TG hearts, consistent with increased activation of mTORC1 (Fig. 4B).

To look for downstream consequences of mTORC1/ISR activation, we measured levels of transcripts that are typically induced (e.g., *Mthfd2* and *Fgf21* (13, 15)). After 10 wks on Doxy diet, there were no differences between WT and TG hearts (SFig. 5). In contrast, TG hearts from 18-wk Doxy-fed mice had higher mRNA levels of one-carbon metabolism enzymes, components of amino acid transport/metabolism, *Fgf21* and *Atf4* (Fig. 4C). There was no difference in *Atf2*, *Atf3, Atf5* and nor in *Gdf15* or in the proteases *Clpp* and *Lonp1* (Fig. 4C). Additionally, protein levels of ATF4, as well as MTHFD2 and ASNS (two ATF4 targets) were substantially elevated in TG heart lysates (Fig. 4D).

**Figure 5:**
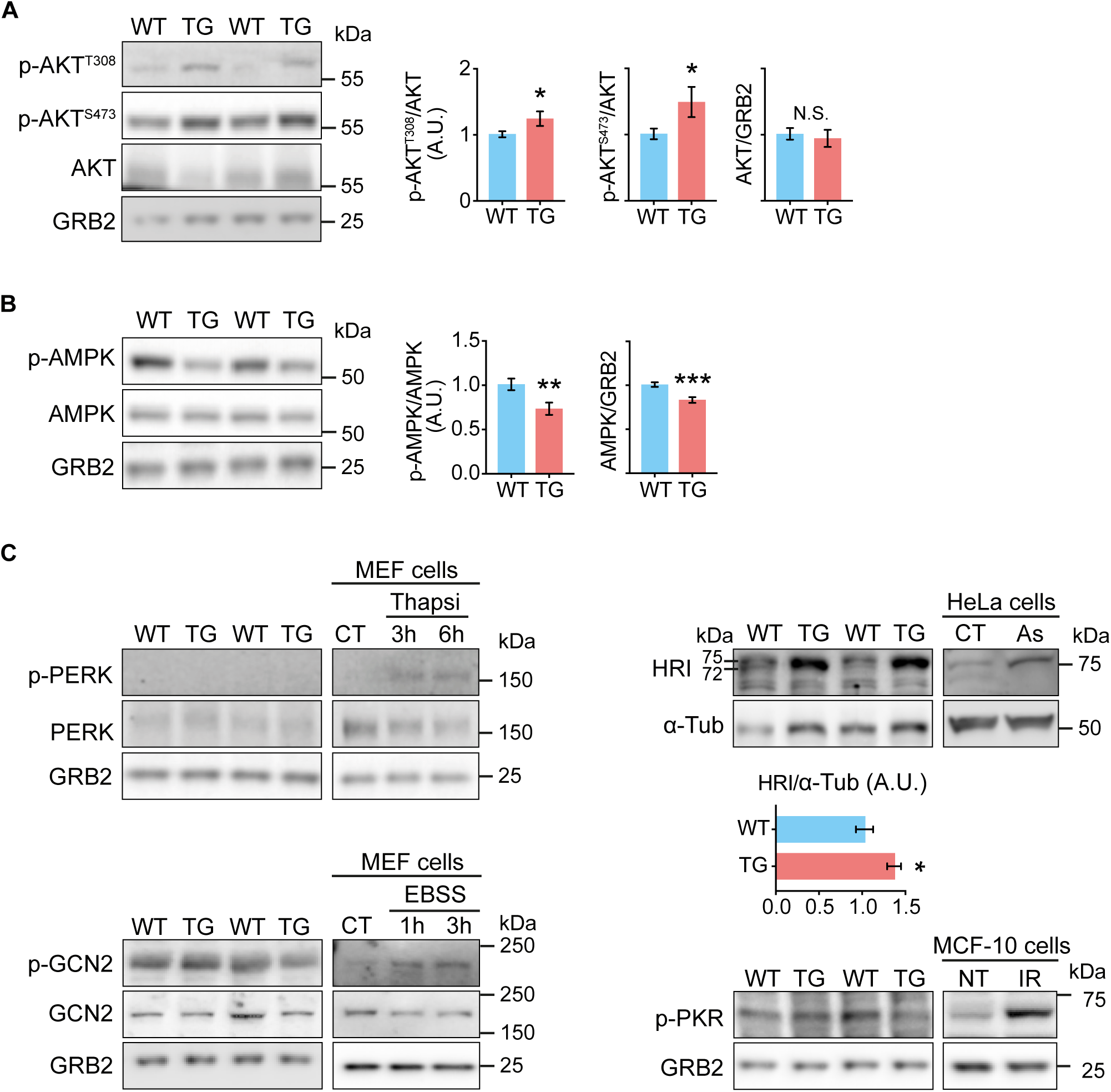
Altered activation of kinases upstream of eIF2α and mTORC1 in hearts depleted of FXN for 18 wks. All measurements were done in heart lysates from WT and TG mice fed Doxy for 18 wks. **(A)** AKT signaling. Representative immunoblots of p-AKT (Ser473), p-AKT (Thr308), total AKT, and GRB2 and GAPDH (loading controls). *Right*: Quantification (n= 9 WT/7 TG). **(B)** AMPK signaling. Representative immunoblots of p-AMPK (Thr302), total AMPK and GRB2 (loading control). *Lower panel*: Averaged p-AMPK/AMPK, p-AMPK/GRB2 and AMPK/GRB2 values (n=9 WT/7 TG). **(C)** Kinases upstream of eIF2α: PERK, GCN2, PKR and HRI. Representative immunoblots of p-PERK (Thr980), total PERK, p-GCN2 and total GCN2, and p-PKR, with GRB2 as loading control. N=6/genotype. Controls for antibodies: p-PERK: mouse embryonic fibroblasts (MEF) treated with thapsigargin (1 µM) to induce ER stress; p-GCN2: MEF cells cultured for 6 hours in EBSS (Earle’s Balanced Salt Solution) to deprive cells of amino acids; p-PKR: MCF-10 cells were irradiated (20gy) to cause DNA damage to activate PKR; HRI: HeLa cells were incubated in 100 µM Na-Arsenite (As) for 1 hour. All values are the mean ± s.e.m. Statistical comparison was by unpaired t-test, \**p*<0.05, \*\**p*<0.01, \*\*\**p*<0.001. N.S.: not significant.

### Insights into upstream regulators of eIF2α and mTORC1

To gain insight into the drivers of the elevated p-eIF2α and mTORC1 signaling in hearts from 18-wk Doxy-fed mice, we assessed the activation of AMPK, a negative regulator of mTORC1, and of AKT, an activator of mTORC1. AKT has also been reported to inhibit AMPK in the heart (42, 43). Phosphorylation of AKT was increased at both phosphorylation sites (Thr308 and Ser473) in TG heart lysates, indicating increased AKT activation (Fig. 5A). We measured the phosphorylation status of AMPK and found a consistent decrease in p-AMPK in FXN-depleted hearts, supporting suppressed AMPK signaling (Fig. 5B).

Four kinases have been identified that phosphorylate eIF2α: PERK, GCN2, PKR and HRI (8). Their activation was evaluated by phosphorylation status, via immunoblotting. p-PERK was undetectable in heart lysates from both WT and TG mice; reliability of the p-PERK antibody was confirmed using mouse embryonic fibroblasts treated with thapsigargin to induce endoplasmic reticulum stress (Fig. 5C). Additionally, BIP/GRP78 levels, usually increased with ER stress, were unchanged (SFig. 6A). Both p-GCN2 and p-PKR were detectable, but levels were similar in WT and TG hearts (Fig. 5C). Reliability of the p-GCN2 and p-PKR antibodies was confirmed using positive controls (see Fig. 5C for a description of the controls). Finally, HRI was evaluated using an antibody that detects total HRI, since an antibody against p-HRI is not available. When samples are run slowly on a 7% gel; activated HRI is taken as the presence of two bands (that can appear merged) with the higher molecular weight band being more intense, or an apparent shift of a more intense band to a higher molecular weight (e.g., (6, 44, 45)). In heart lysates, we detected a strong band at ∼75 kDa in TG samples, whereas a less intense doublet was detected in WT samples (Fig. 5C, and see also SFig. 6B). As a positive control for the HRI antibody, HeLa cells were treated with arsenite (e.g., (44)); a single band that increased in intensity with arsenite was evident (Fig. 5C).

Finally, because mTORC1 and eIF2α activation have opposite implications for protein synthesis, we evaluated global protein translation *in vivo*, using the SUnSET method that evaluates puromycin incorporation into elongating peptide chains (46). A decrease in the signal from the anti-puromycin antibody in heart lysates from 18-wk Doxy-fed TG mice indicated that global protein synthesis was lower in FXN-depleted hearts (Fig. 6A). To have a more complete picture of proteostasis in the hearts from 18-wk Doxy-fed mice, insight into protein breakdown was obtained. Transcript levels of E3 ubiquitin ligases, Atrogin1 (*Fbxo32*) and MuRF1 (*Trim63*), were similar between WT and TG hearts (Fig. 6B). Consistent with a lack of difference in degradation, the amount of polyubiquitinated proteins with specific lysine (K) linkages (K48 and K63) was similar between WT and FXN-depleted hearts (Fig. 6C). Finally, insight into autophagy was obtained by evaluating protein levels of LC3II, the lipidated form of LC3I, and p62. LC3II relative to LC3I was similar between WT and TG hearts, whereas p62 expression was lower, suggesting that autophagy was not elevated in TG hearts from 18-wk Doxy fed mice.

## DISCUSSION

The main findings and proposed mechanistic links are shown in Fig. 6D. After 18 wks on Doxy diet, and at least 8 wks with FXN levels < 20% of normal, TG hearts did not display overt hypertrophy and were in fact smaller. Cardiac contractility was maintained, likely due to maintained β-oxidation, though oxphos capacity for pyruvate was lower. The bioenergetics changes generally matched the proteomics changes and were not entirely linked to a loss of ISC proteins. Global protein translation was lower, while protein degradative pathways were unaltered. In parallel, there were perturbations in the main nutrient signaling pathways, AMPK and mTORC1, and in AKT, a regulator of those pathways. Activated AKT could lead to less AMPK phosphorylation and drive mTORC1 signaling. Lower AMPK activity would also be permissive for mTORC1 activation. The ISR was also activated, likely by HRI. Phosphorylation of eIF2α would lower global protein translation, possibly explaining the lack of hypertrophy. We propose that the response to FXN loss in this model at least partially reflects ISR activation that predominates over mTORC1 signaling, in turn lowering the ATP demanding process of protein translation, leading to a smaller heart weight that helps to sustain lower ATP demand. We speculate that this response is protective for the FXN-depleted heart, at least in the short term. We further propose that the 18-wk Doxy-fed mice represents a pre-clinical model of FRDA.

### The TG model

The mouse model used here is the same genetic model as that used by Chandran (27). However, different Doxy regimens were used. Chandran supplied Doxy by intraperitoneal injection and in drinking water whereas we supplied Doxy in the chow. The time course and extent of FXN loss in the heart, and the body weight loss, were similar for the two Doxy regimens. Yet, mortality of the mice used in the present study was lower; our mice showed ∼35% mortality by 18 wks of Doxy whereas Chandran observed ∼35% mortality by 10 wks and >50% by 18 wks of Doxy. Chandran tested several Doxy doses; higher dose associated with greater mortality. We propose that the dose used in the present study was lower and/or less toxic than the lowest dose used by Chandran (and on which they reported phenotypes).

### Heart size and function

After 18 wks of FXN depletion, heart weight relative to tibial length was lower and cardiac fiber cross-sectional area was slightly but significantly smaller. There may have been some preservation of LV mass, based on LV wall and interventricular septal wall thicknesses that were similar between TG and WT. Nonetheless, it is clear that TG hearts did not display overt hypertrophy observed in MCK mice (6, 22–24). The TG mice studied by Chandran, under the different Doxy dosing regimen, showed evidence for hypertrophy after 24 wks of Doxy, but not after 12 wks (27); an intermediate time point was not investigated. Often cardiac stress is associated with hypertrophy. Yet, decreased heart mass has been documented in mice in the context of fasting (47) (48, 49), and dexamethasone (50) and short term doxorubicin treatments (51). Lower heart mass with fasting, including prolonged fasting, was associated with elevated p-AMPK (48) and LC3-II (47) (48, 49), and lower p-mTORC1 (48), the opposite of what was observed in TG hearts. With doxorubicin and dexamethasone, heart weight decrease was Trim63 (MuRF-1)-dependent (50, 51), whereas lower TG heart weight was not associated with increased protein degradation. Thus, the mechanisms underlying the lower TG heart weight appear distinct from those responsible for doxorubicin, dexamethasone and starvation. We propose that TG heart weight reflects normal protein degradation in conjunction with lower global protein synthesis.

After 18 wks on Doxy, hearts from TG mice showed no sign of fibrosis and had maintained contractility. This differs from the MCK model in which cardiac fibrosis, lower ejection fraction and markers of apoptosis were evident (6, 21–24). Chandran observed lower ejection fraction and fibrosis after 24 wks of Doxy, but not after 12 wks (27). Intermediates time points were not evaluated. We performed echocardiography on a small number of TG mice after 20 wks of Doxy diet; ejection fraction continued to be normal, though there was evidence for hypertrophy (not shown). We decided to not pursue time later time because weight rapidly became significant, mice felt cold, and mortality increased.

Cardiac contractility was maintained and changes in transcript levels of glycolytic and β-oxidation enzymes that are associated with a cardiac energy crisis (37) were absent. These findings suggest that cardiac energy production was sufficient in TG hearts. The most likely explanation for this is that the heart preferentially oxidizes fatty acids and β-oxidation was relatively preserved in TG heart mitochondria. Greater oxphos efficiency, evidenced by lower proton leak, might also be a factor. Another consideration is that, though maximal oxphos capacity for pyruvate was lower, this maximal capacity includes a reserve, such that sufficient capacity was still available for the pyruvate contribution to ATP production. Finally, the higher PDK4 in TG hearts may allow for a further increase in the use of fatty acids over glucose for ATP production.

### Mitochondrial bioenergetics and proteomics

A detailed bioenergetics analysis was performed in FXN-depleted heart mitochondria, which was lacking in other models of FRDA. Our analysis showed the expected decrease in oxphos capacity. However this was substrate-dependent such that β-oxidation was relatively preserved. One contributor to preserved β-oxidation could be that electrons from acyl-CoA dehydrogenase-coupled reduction of FAD^+^ enter Complex III via ubiquinol (see Fig. 2A). Thus, β-oxidation, differently from pyruvate oxidation, does not depend for reducing equivalent generation on the TCA cycle and Complex I, the pathways with the greatest decrease in protein abundance in TG hearts. Electron flow from FADH2 into Complex III depends on ETFDH, an ISC-containing protein whose abundance was lower, as was abundance of short-chain specific acyl-CoA dehydrogenase. However the acyl-CoA dehydrogenases have overlap in substrate specificity, and the rest of the β-oxidation pathway, and Complex III, IV and ATP synthase were minimally changed with FXN loss. This contrasts with the TCA cycle and Complex I for which 6/14 and 15/36 proteins, respectively, were significantly downregulated. Thus, in the TG hearts, pyruvate oxidation would be more likely than β-oxidation to encounter bottlenecks.

The heart mitochondrial proteome presented here, from 18-wk Doxy-fed mice, is, to our knowledge, the first broad proteomics analysis of FXN-depleted mitochondria. The dataset provides novel insights, and also shows similarities with a proteomics analysis of heart mitochondria from mouse models of mtDNA depletion (39). In particular, Kuhl (39) found broad decreases in oxphos proteins, as we did, as well as decreases in ubiquinone biosynthetic enzymes (also found in our data: Supplemental Table). Kuhl reported upregulation of CYB5R1, of enzymes promoting conversion of glutamate to proline (PYC1 and PYC2), the serine protease HTRA2 and caspase activator DIABLO, all of which we found (Supplemental Table). A prominent difference in our proteomics data is the absence of change, or decrease, in heat shock proteins (HSPD1, HSPE1, HSPA9) and the proteases CLPP and LONP1; all the latter were strongly increased in mtDNA-depleted hearts (39). We also found PMPCA and PMPCB to be substantially lower; Kuhl did not report protein levels but found no change in PMPCB mRNA levels. Thus, the proteomics data point to similarities between FXN-depleted and mtDNA-depleted hearts; indeed, mtDNA depleted hearts showed activated ISR (39). Differences between FXN and mtDNA depletion in how protease abundance is altered suggests that mitochondrial proteases are responsive to a mechanism other than the ISR, and that differs between the two stress conditions.

Levels of ISC-containing proteins were expected to be lower (e.g. (23)). Indeed, FECH was substantially lower, and three Complex I subunits, SDHB and ETFDH were significantly, though modestly decreased. Yet the other ISC-, heme and lipoic-acid containing proteins were not significantly lower. Immunoblot analysis of several proteins in heart lysates and isolated mitochondria were confirmatory. A broad survey of the literature shows that ISC-containing proteins and their activity are not unequivocally and uniformly decreased when ISC biosynthetic machinery is lost. Heart mitochondria from the MCK mouse, in which FXN is fully lost from muscle soon after birth, showed lower SDHB protein only when mice were older than 4 wks of age (23), whereas FECH was substantially lower by 3 wks and barely detectable by 5 wks (6). In fibroblasts lacking ISCU, FECH was barely detectable whereas aconitase was still expressed, though less so (52). When enzyme activity is considered, Complex I activity was more sensitive to loss of FXN or other ISC biosynthetic components than SDH or Complex III activity (53–56). In MCK mouse hearts, SDH activity was more sensitive to FXN loss than aconitase activity (21). To our knowledge this is the first discussion of a non-uniform effect of FXN loss on abundance and activity of ISC-containing proteins. We cannot explain this phenomenon, though mechanisms can be proposed. Longer average half-life for Complex III and IV subunits compared to Complex I and SDH subunits (Supplementary Data 1 in (57)) suggests differential protein stability as a factor. Altered abundance of mitochondrial proteases could be another factor. Matrix-localized CLPP was significantly less in TG hearts (see Supplemental Table); the N module of Complex I is a substrate for CLPP (58), thus CLPP downregulation may have actually mitigated the loss of Complex I in TG hearts. Also, PO2 of cardiac mitochondria was estimated to reach levels as low as 10 mmHg (59), raising the question of whether FXN-independent ISC biogenesis shown to occur in hypoxia (60) is possible in cardiac mitochondria.

### Nutrient and stress signaling

Overt cardiac pathology was not apparent in 18-wk Doxy-fed TG mice. Yet the lower heart weight, and substantial literature describing changes in nutrient and stress signaling associated with mitochondrial disease (10-15, 39, 61), suggested that nutrient and stress signaling might be altered in TG hearts. Moreover, MTHFD1L, barely detectable in WT mitochondria, was readily detected in TG and is an ATF4 target (38). The upregulation of other ATF4 targets and ATF4 itself that we documented suggested that the ISR and/or mTORC1 was activated. ISR activation and upregulation of ATF4 targets were reported in the heart of MCK FXN mice (6), though mTORC1 signaling was not evaluated. We found that both the ISR and mTORC1 were activated in the heart of 18-wk Doxy TG mice. Elevated p-AKT and lower p-AMPK could both have contributed to mTORC1 activation (62). Elevated p-AKT has also been linked to increased intracellular iron (63), and was evident in TG hearts. The lower p-AMPK was surprising, but has been reported in hearts depleted of acyl-CoA synthase-1 (64). Moreover, higher p-AKT in the heart was associated with lower p-AMPK (42, 43), likely via phosphorylation of the AMPK-α1 subunit which prevents its activation (65).

Of the four kinases shown to phosphorylate eIF2α, only HRI showed evidence of activation in TG hearts. Evidence for HRI activation was also found in the MCK mouse heart, though activation was not sustained whereas elevated p-eIF2α was (6); phosphorylation status of the other kinases was not evaluated in MCK mouse hearts. Heme depletion leads to HRI activation (66), and may have occurred in TG (and MCK) hearts because FECH, required for heme synthesis, was strongly decreased. HRI can also be activated by DELE1 upon its cleavage and release of a soluble fragment from the inner mitochondrial membrane (67, 68). DELE1 was not detected in our proteomics data nor in transcriptomics data reported by Chandran (27).

Elevated mTORC1 signaling would predict hypertrophy (e.g. (64)). Yet, overt hypertrophy was not evident in TG hearts, and hypothesize that this reflects lower global translation due to higher p-eIF2α contributed. MCK mice also showed elevated p-eIF2α (6), however cardiac hypertrophy was substantial (6, 21, 23). Presence of hypertrophy in MCK mice suggests that mTORC1 signaling was elevated and predominated over the ISR; mTORC1 signaling was not evaluated in MCK mice. Interestingly, in the later study, total eIF2α increased (Fig. 4A in (6)); thus, though eIF2α phosphorylation also rose in MCK hearts, the amount of non-phosphorylated eIF2α may have been maintained, allowing for increased global protein translation and hypertrophy. There was no sign of higher eIF2α in TG hearts. Our study demonstrates that elevated mTORC1 signaling and the ISR can coexist and shows the benefit of evaluating both pathways. We propose that the extent of cardiac hypertrophy can depend on both the ISR and mTORC1.

### Phenotype *vs*. FXN level

FXN abundance at ∼20% of the normal level was sufficient to preserve bioenergetics parameters (though pyruvate oxidation trended lower), was not associated with changes in expression of ATF4 target genes, nor in FECH protein abundance. Thus, as little as 20% of normal FXN levels may be enough to prevent pathology. Yet, a few points should be considered. Very mild impairment in bioenergetics has been associated with eIF2α phosphorylation (69), and alterations in signaling in response to mitochondrial dysfunction can take time to become manifest (15). Furthermore, the bioenergetics profile at the 10-wk Doxy time point may have been influenced by protein stability; thus it would be interesting to determine how the heart would respond to a sustained FXN level of ∼20% of the normal level. It should also be noted that, while FXN protein is undetectable in TG hearts after 18 wks of Doxy, substantial oxphos capacity and ETC capacity remained. Even FECH levels, though substantially decreased, were 50% the WT level, and levels of LIAS were normal. Thus, discordance between FXN abundance and phenotypes, possibly related to protein stability and/or time-dependence of yet-to-be-understood cellular changes, needs to be considered here and likely in other FRDA models.

### Implications

We show that several changes in nutrient and stress signaling can occur in the FXN-depleted heart. The relative extent of these changes might differ among individuals with FRDA and contribute to the variability in cardiac phenotypes, such as presence or absence of hypertrophy. We also show that the impact of FXN loss on substrate metabolism can be substrate-dependent, raising the question of whether vulnerability to FXN depletion can depend on basic metabolic phenotype; in particular, neurons have high ATP demand and primarily rely on aerobic glucose oxidation with little capacity for β-oxidation. Our study also indicates that normal heart function, energy production and lack of hypertrophy can be misleading indicators that the FXN-depleted heart is normal. Finally, small molecules exist for the AMPK, mTORC1 and ISR pathways, raising the question of whether these pathways could be targeted to treat FRDA, as suggested for other mitochondrial diseases (12, 17, 20). However, a better understanding of the impact of the changes in these pathways on organ function in the FXN-depleted heart will first need to be understood.

## MATERIALS and METHODS

### Mouse model

Mice were used in accordance with mandated standards of care and, use was approved by the Thomas Jefferson University Institutional Animal Care and Use Committee. Mice were housed at 22C, under a 12-hr light-dark cycle (lights on at 07:00). We used a doxycycline-inducible model of FXN knockdown (27). Here, TG mouse (on C56BL/6J background) contain a Tet-On frataxin shRNA expression cassette. Littermates that do not contain the shRNA cassette were used as controls (WT). Doxycycline was administered in the chow (200 p.p.m. in Chow 5SHA, Animal Specialties) to WT and TG mice starting at 9 weeks of age. Only male mice were used for experiments.

### Cell culture and stress conditions for antibody positive controls

MEF and HeLa cells were cultured in Dulbecco’s modified Eagle’s medium (11965-118, Invitrogen) supplemented with 10% FBS (fetal bovine serum), 2 mM glutamine and 100 U/ml penicillin and 100mg/ml streptomycin and maintained at 37°C in a 5% CO2-humidified air. A series of positive controls were generated using stress conditions to validate the activation of the ISR kinases (PERK, GCN2, HRI and PKR) by their respective antibodies. For p-PERK antibody, MEF cells were treated with thapsigargin 1μM for 3 and 6 h, to have a positive control for ER stress. For p-GCN2 antibody, MEF cells were cultured in EBSS (5.33 mM KCl, 26.19 mM NaHCO3, 117.24 mM NaCl, 1.104 mM NaH2PO4 and 5.56 mM D-Glucose) for 3 and 6 h, to generate an amino acid deprivation condition. For HRI activation, HeLa cells were treated with arsenite (100, 200 and 300 μM AsNaO2) for 1 h. Finally, a positive control for p-PKR antibody was generated using MCF-10 cells irradiated with a CellRad350 at 0.8-0.9 Grey per minute.

### Echocardiography

Transthoracic two-dimensional M-mode echocardiography was performed using an echocardiographic imaging system (Vevo 2100, VisualSonic, Toronto, Canada) with a 40-MHz probe. Mice were anesthetized through the inhalation of isoflurane (∼1–2%); anesthesia was titrated with the goal of equalizing heart rate in all mice. M-mode interrogation was performed in the parasternal short-axis view at the level of the greatest left ventricular end-diastolic dimension. Left ventricular wall thickness and internal dimensions were measured and used for calculation of fraction shortening and ejection fraction values.

### Isolation of heart mitochondria

Heart mitochondria were isolated as described in (60). All steps were performed on ice or at 4°C. The heart was dissected, washed in heart isolation buffer (HB: 225 mM mannitol, 75 mM sucrose, 20 mM HEPES, 0.1 mM EGTA, pH 7.4), minced with a razor blade, then suspended in HB+0.5% defatted BSA in a glass/Teflon Potter-Elvehjem homogenizer and homogenized (350 rpm, 10 passes). Samples were centrifuged (5 min, 500*g*), the supernatant was centrifuged (10 min, 9000*g*), then the pellet was resuspended in HB devoid of BSA then centrifuged again (10 min, 9000*g*). The final pellet was resuspended in HB (no BSA) in a volume that resulted in a protein concentration of ∼20 mg/ml. Protein concentration was determined by bicinchoninic acid (BCA) assay (ThermoFisher Scientific 23228, 1859078).

### Bioenergetics analyses in isolated heart mitochondria

For most experiments, O2 consumption (JO2) was measured using the Seahorse XF24 Analyzer (Seahorse Bioscience, Billerica, MA, USA). Isolated mitochondria were studied essentially as we have done previously (70). Each well of the custom microplate contained 10 µg of mitochondria suspend in mitochondria assay medium (MAS; 70 mM sucrose, 22 mM mannitol, 10 mM KH2PO4, 5 mM MgCl2, 2 mM Hepes, 1 mM EGTA, 0.2% defatted BSA, pH 7.4 at 37°C). Amount of mitochondria had been optimized such that the O2 *vs*. time signal was linear under all conditions. The microplate was centrifuged (2000*g*, 20 min, 4°C) to promote adhesion of mitochondria to the plastic. Attachment was verified after centrifugation and again after experiments. Different substrates were tested: malate-pyruvate (5 mM/10 mM), succinate (10 mM + 1 µM rotenone to inhibit Complex I and thereby prevent reverse electron flow through that complex), palmitoyl-L-carnitine – malate (20 µM/1 mM), octanoyl-L-carnitine (200 µM + 1 mM malate). Oligomycin (4 µg/mL) was used to measure non-phosphorylation “leak” respiration JO2 and the uncoupler FCCP (6.7 µM) was used to measure maximal electron transport chain activity. Antimycin titrations of octanoyl-L-carnitine (200 µM + 1 mM malate) were done in the presence of oligomycin (4 µg/ml) using the Oroboros O2k, with 200 µg of mitochondria per reaction, in MAS.

### Fluorometric measurements in isolated mitochondria

Fluorometric measurements of mitochondrial membrane potential (ΔΨm) were performed using a multiwavelength-excitation dual-wavelength-emission fluorimeter (DeltaRAM, PTI). Briefly, isolated mitochondria (375 µg) were resuspended in 1.5 ml of intracellular medium containing 120 mM KCl, 10 mM NaCl, 1 mM KH2PO4, 20 mM Tris-HEPES at pH 7.2 and maintained in a stirred thermostated cuvette at 36 °C. TMRM (1μM) was added prior the experiment. Antimycin titrations of octanoyl-L-carnitine (200 µM µM + 1 mM malate) were done in the presence of oligomycin (4 µg/ml). FCCP (6.7 μM) was used to obtain maximal membrane depolarization.

### Immunoblot analysis

For western blot analysis, heart was quickly frozen in liquid nitrogen. After that, tissue was homogenized with a lysis buffer containing: 150 mM NaCl, 25 mM HEPES, 2.5 mM EGTA, 1% Triton 100X, 1% Igepal (10%), 0.10% SDS, 0.1% Deoxycholate, 10% Glycerol, protease inhibitor (Roche 11873580001) and phosphatase inhibitor cocktail (200mM sodium fluoride, 200mM, imidazole, 115mM sodium molybdate, 200mM sodium orthovanadate, 400mM sodium tartrate dihydrate, 100mM sodium pyrophosphate and 100mM β-glycerophosphate), using a glass/Teflon homogenizer at 300 rpm. After that, heart lysates were incubated at 4°C for 45 min, then centrifuged at 18000*g* for 20 min. From the supernatant, protein concentration was measured by BCA assay. Specifically, for LC3B detection in heart, we used T-PER tissue protein extraction reagent (ThermoFisher Scientific 78510). Hearts were quickly pulverized over dry ice, and then incubated for 1h in T-PER at 4°C (shaking them at 1500 r.p.m.). After that, samples were centrifuged at 18000*g* for 30 min at 4°C. From the supernatant, protein concentration was measured by BCA assay. For western blot analysis from cultured cells, cell lysis was obtained using RIPA buffer containing protease and phosphatase inhibitors. Cell lysates were spun at 12000 rpm for 10 min. From the supernatant, protein concentration was measured by BCA assay. For western blot analysis of mitochondrial proteins, isolated heart mitochondria were resuspended in 4X sample buffer. Primary antibodies were used for overnight incubation diluted in TBS-T 1% and are listed below.

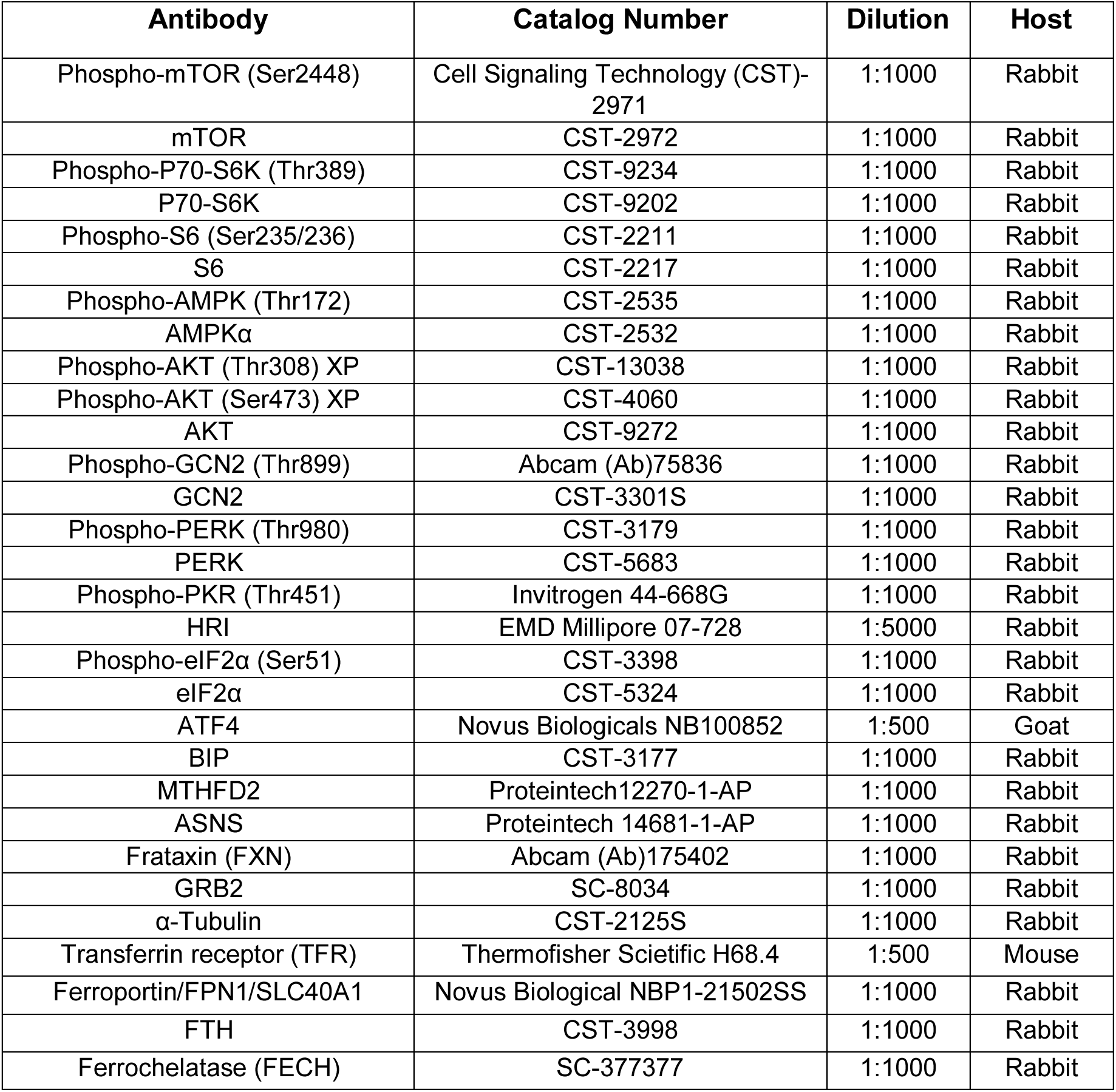
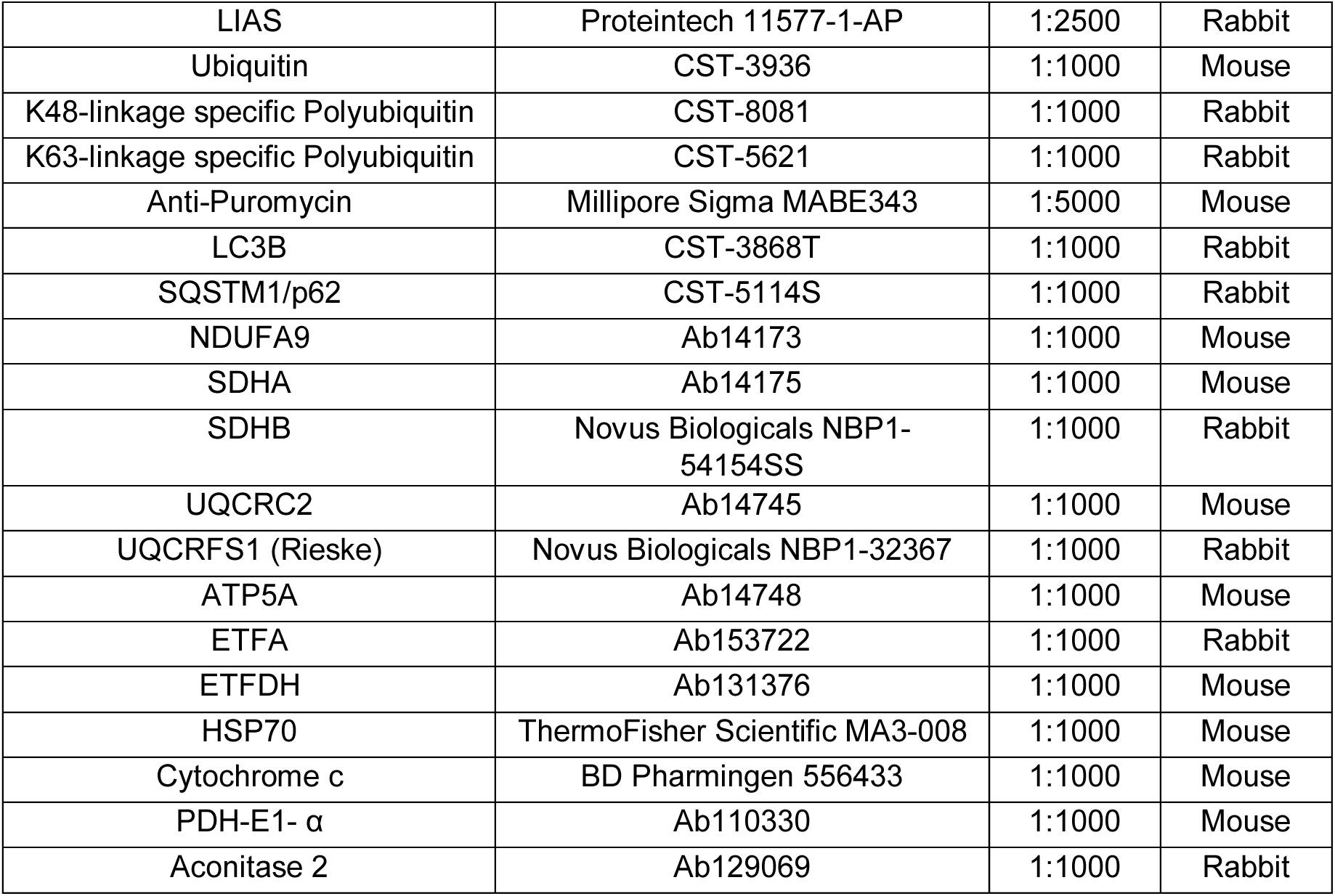
Primary Antibodies used in immunoblot analysis

### Blue native electrophoresis and in-gel activity

Isolated heart mitochondria (100 µg) were lysed in 4% digitonin in extraction n buffer (30 mM HEPES, 12 % glycerol, 150 mM potassium acetate, 2 mM aminocaproic acid, 2 mM EDTA disodium salt, protease inhibitor tablet (Roche 11873580001), pH 7.4), at 4°C for 30 min with constant shaking at 1500 rpm. The samples were centrifuged at 25000 g, 20 min, 4°C, then 1:400 diluted 5% Coomassie Blue G-250 was added to the supernatant. The samples were then loaded on a gradient gel (NativePAGE™ NovexR 3– 12% Bis-Tris). The gel was run overnight in Native PAGE buffers (Invitrogen). The gel for Coomassie blue staining was run in dark blue buffer until the dye front reached 1/3 of the gel (150V) then in light blue buffer for the rest of the time up to 16h (30V). For gels used for in-gel activity, the gel was run first in light blue buffer then in clear buffer (Native PAGE Invitrogen). Following the electrophoresis, the gel was stained with Coomassie Blue R-250 for 30 min, then destained (40% MetOH, 10% acetic acid in water). In-gel activity of complex I, II, and V was run as described (71).

### Proteomics: LC-MS/MS Analyses and Data Processing

Samples (25 µg each) were run into a NuPAGE 10% Bis-Tris gel (Thermo Scientific) for a short distance, and the entire gel lane was excised and digested with trypsin. Liquid chromatography tandem mass spectrometry (LC-MS/MS) analysis was performed using a Q Exactive HF mass spectrometer (Thermo Scientific) coupled with an UltiMate 3000 nano UPLC system (Thermo Scientific). Samples were injected onto a PepMap100 trap column (0.3 x 5 mm packed with 5 μm C18 resin; Thermo Scientific), and peptides were separated by reversed phase HPLC on a BEH C18 nanocapillary analytical column (75 μm i.d. x 25 cm, 1.7 μm particle size; Waters) using a 4-h gradient formed by solvent A (0.1% formic acid in water) and solvent B (0.1% formic acid in acetonitrile). Eluted peptides were analyzed by the mass spectrometer set to repetitively scan m/z from 400 to 1800 in positive ion mode. The full MS scan was collected at 60,000 resolution followed by data-dependent MS/MS scans at 15,000 resolution on the 20 most abundant ions exceeding a minimum threshold of 20,000. Peptide match was set as preferred, exclude isotope option and charge-state screening were enabled to reject unassigned and single charged ions.

Peptide sequences were identified using MaxQuant 1.6.17.0 (72). MS/MS spectra were searched against a UniProt mouse protein database (October 2020) and a common contaminants database using full tryptic specificity with up to two missed cleavages, static carboxamidomethylation of Cys, and variable Met oxidation, protein N-terminal acetylation and Asn deamidation. “Match between runs” feature was used to help transfer identifications across experiments to minimize missing values. Consensus identification lists were generated with false discovery rates set at 1% for protein and peptide identifications. Statistical analyses were performed using Perseus 1.6.14.0 (73). Missing values were imputed with a minimum value, and t-test p-values were adjusted (q-value) to account for multiple testing using Benjamini-Hochberg FDR function in Perseus.

The mitochondrial proteins were selected by matching gene names with a database of known and predicted mitochondrial genes using Python : the ’Mitochondrial Part’ annotation from the Jensen Compartments database [https://doi.org/10.1093/database/bau012] downloaded from Enrichr (https://maayanlab.cloud/Enrichr/).

### Quantitative polymerase chain reaction

Total RNA was extracted from tissues using Trizol® (Invitrogen, Carlsbad, CA) then treated with RQ1 DNase (Promega, Madison, WI, USA) at 37°C for 30 mins. RNA concentration was measured by Qubit® Fluorometer (Invitrogen, Carlsbad, CA). RNA was reverse transcribed using oligo(dT)20 primers and SuperScript III (Invitrogen, Carlsbad, CA). Primers were designed using Eurofins Primer Design Tool, and checked for specificity and efficiency. Primers sequences are shown below. qPCR reactions were performed using ITaq SYBR green Supermix with ROX (BIO-RAD, Hercules, CA), with 20 ng cDNA/reactions, using an Eppendorf Mastercycler® ep realplex. The ΔΔCt method was used to calculate mRNA levels relative to *Actb* (β-actin). Primers sequences used are listed below.

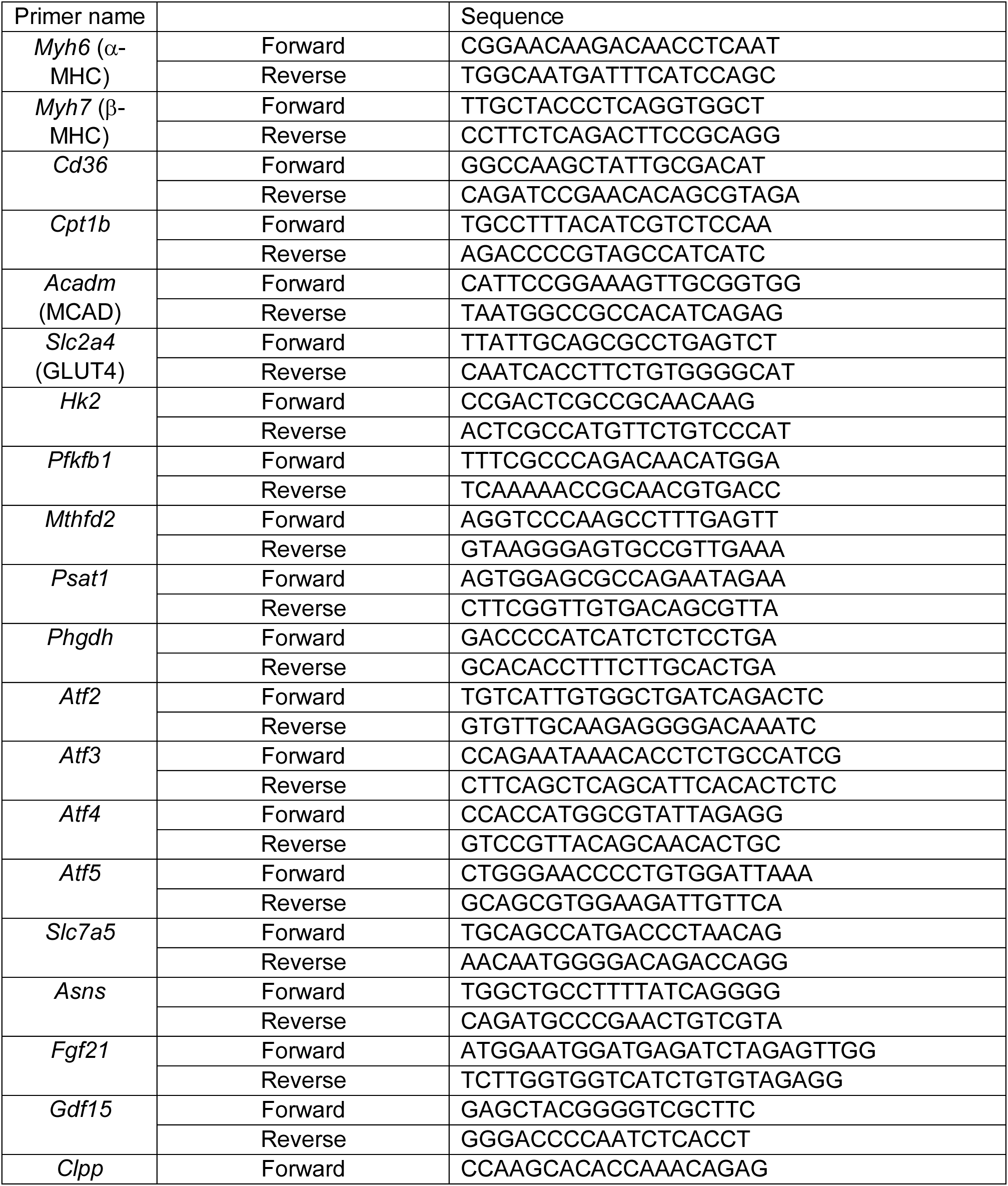
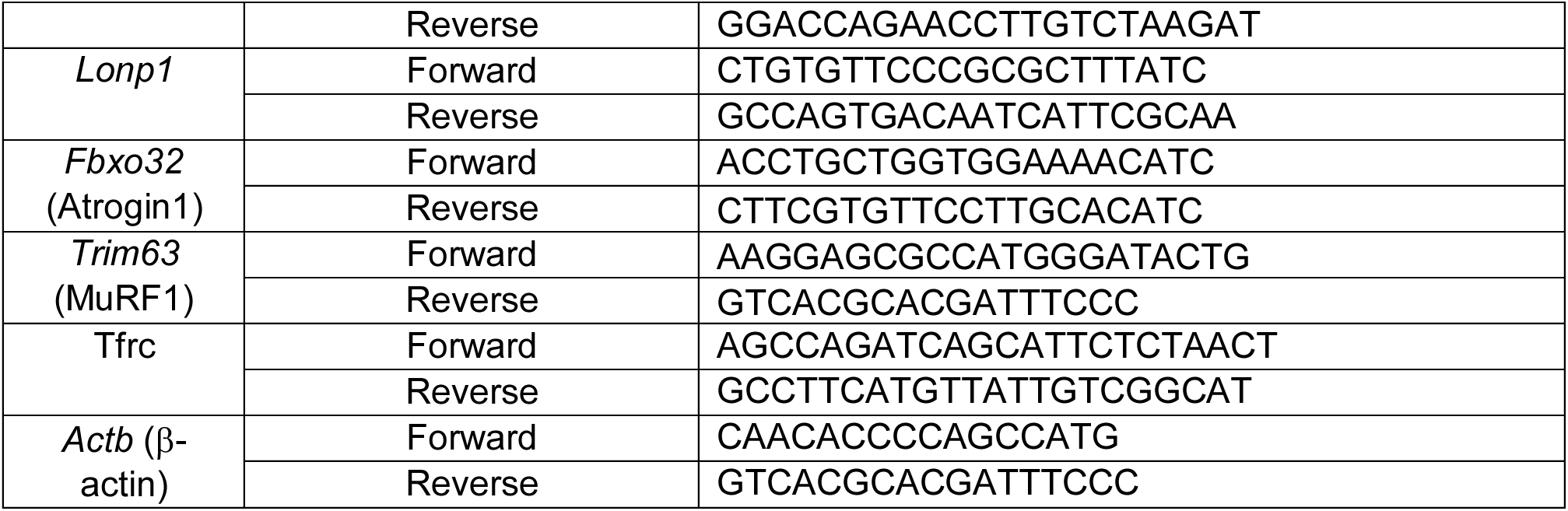
Primers for qPCR (all are mouse-specific)

### Global protein synthesis

To measure the rate of global protein synthesis, we used an in vivo SUnSET assay (46). For this study, a sterilized puromycin (Sigma, 8833) solution in PBS (4mg/mL), was injected intraperitoneally. The volume of the puromycin injection was calculated based on the body weight of the mouse, for a final concentration of 0.04 μmol/g. Thirty minutes after the injection, animals were sacrificed to collect tissue samples for biochemical detection of incorporated puromycin by western blot.

### Histology

Hematoxylin/eosin (H&E), Masson’s Trichrome and Perls’ Prussian Blue staining were performed at the translational research /pathology shared resource at Thomas Jefferson University. Bright field images were acquired using an Olympus CKX41 inverted microscope and cellSens Standard software. Images were analyzed using Image J.

### Quantification and statistical analysis

Data analysis was performed using GraphPad Prism 8.2.1 Software. Unpaired Student’s *t* test for comparison of two means, or one-way analysis of variance followed by Tukey post-hoc comparisons, for multiple comparisons, were performed as appropriate. P value < 0.05 was considered significant. Analysis of the proteomics data is described in the relevant section of Methods, above. In all cases, data not indicated as significant should be considered not statistically different. Details specific for a given measurement, including sample sizes, are provided in Results and the figure legends.

## Supporting information

Supplemental data (Proteomics)

## ACKNOWLEDGEMENTS

The authors thank Drs. Vijay Chandran (University of Florida) and Daniel Geschwind (U.C.L.A.) for providing the mouse model, David Weaver (MitoCare, Thomas Jefferson University) for assistance with proteomics analysis, and Marco Tigano (NYU) for the irradiated MCF-10 cells. Studies were funded by grants from the Friedreich’s Ataxia Research Alliance (to E.L.S.), National Institutes of Health R01 GM123771 (to E.L.S.), National Institutes of Health P30 CA010815 (to Wistar Institute Proteomics & Metabolomics Facility) and National Institutes of Health R50 CA221838 (to H.-Y.T).

---

**Figure S1:**
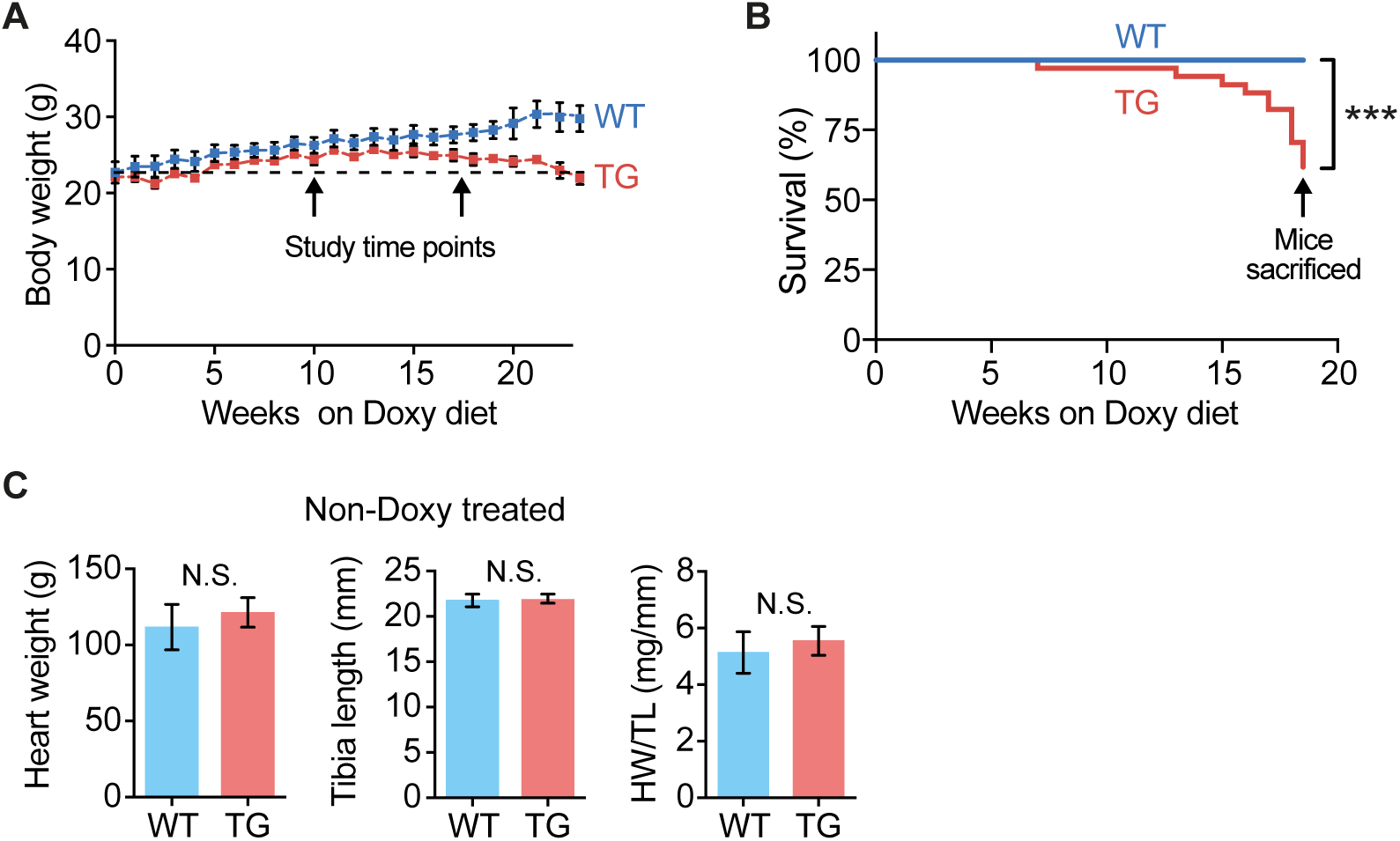
Additional characteristics of the TG mouse model. **(A)** Body weight of WT and TG mice over the course of Doxy feeding. N = 5-12 mice/time point/genotype. **(B)** Survival rate, n=29 WT/34 TG mice, \*\*\**p*<0.001: Curve comparison, Gehan-Breslow-Wilcoxon test. **(C)** No difference in heart weight of WT and TG mice not fed Doxy diet; n = 9 WT/4 TG mice.

**Figure S2:**
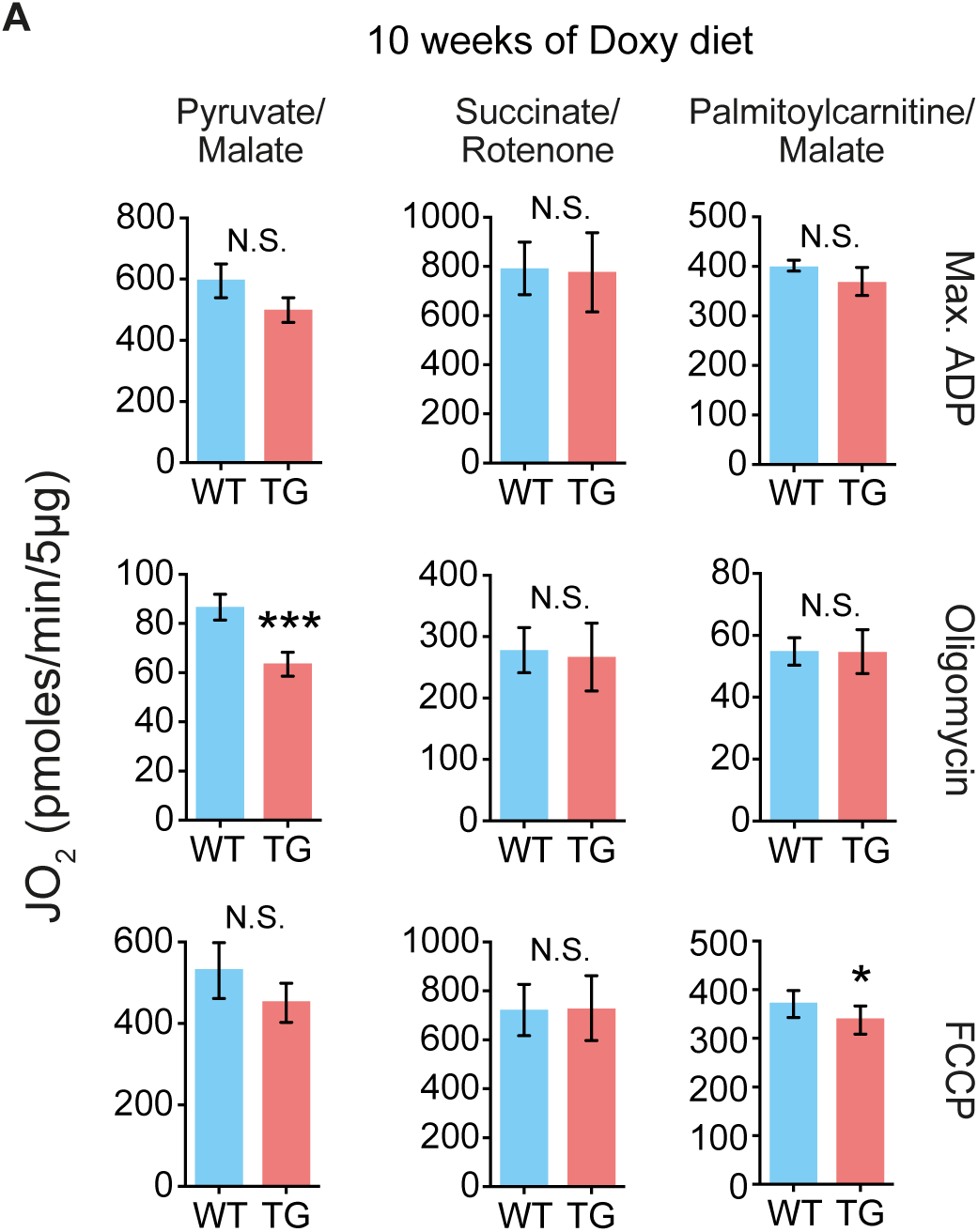
Little effect of FXN depletion on substrate oxidation rates in heart mitochondria from mice on Doxy diet for 10 wks. Oxygen consumption rate (JO2) measured in isolated heart mitochondria supplied with pyruvate/malate (10 mM/5 mM), succinate (10 mM + 1 µM rotenone to prevent electron backflow through complex I), palmitoyl-L-carnitine plus malate (20 µM + 1 mM), or octanoyl-L-carnitine plus malate (200 µM+1 mM). Max. ADP, saturating [ADP] (4 mM), was used to evaluate JO2 reflecting maximal oxidative phosphorylation. Oligomycin was used to evaluated JO2 that reflects maximal leak-dependent oxidation. The chemical uncoupler, FCCP (1 µM), 2.5 µg/ml) as used to evaluate JO2 that reflects maximal electron transport chain capacity under the prevailing substrate conditions. All values represent the mean ± s.e.m. Statistical analysis was by unpaired t-test; *p<0.05, n=6/genotype. Excel file: Mass spectrometry proteomics data set from heart mitochondria isolated from 18-wk Doxy-fed mice

**Figure S3:**
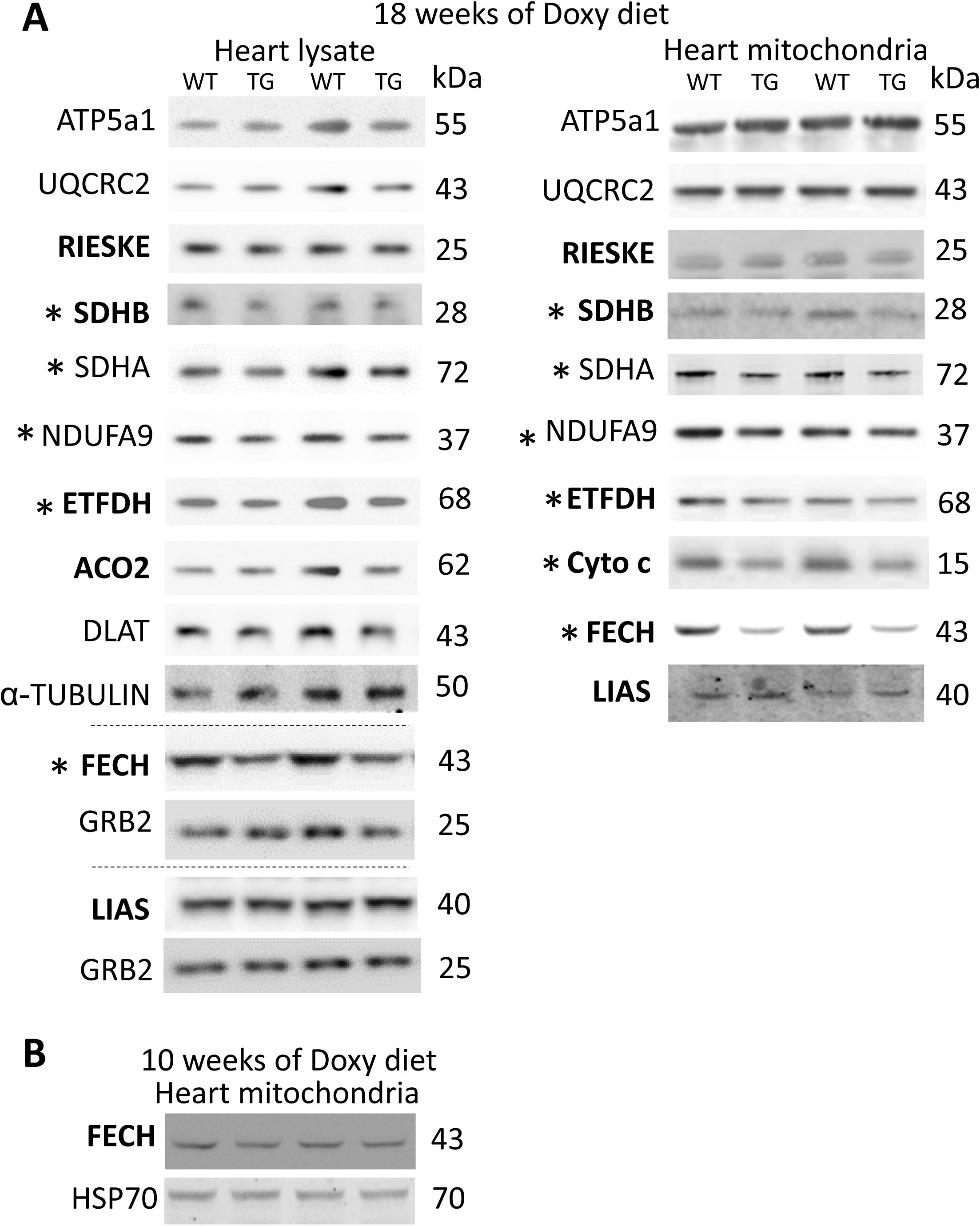
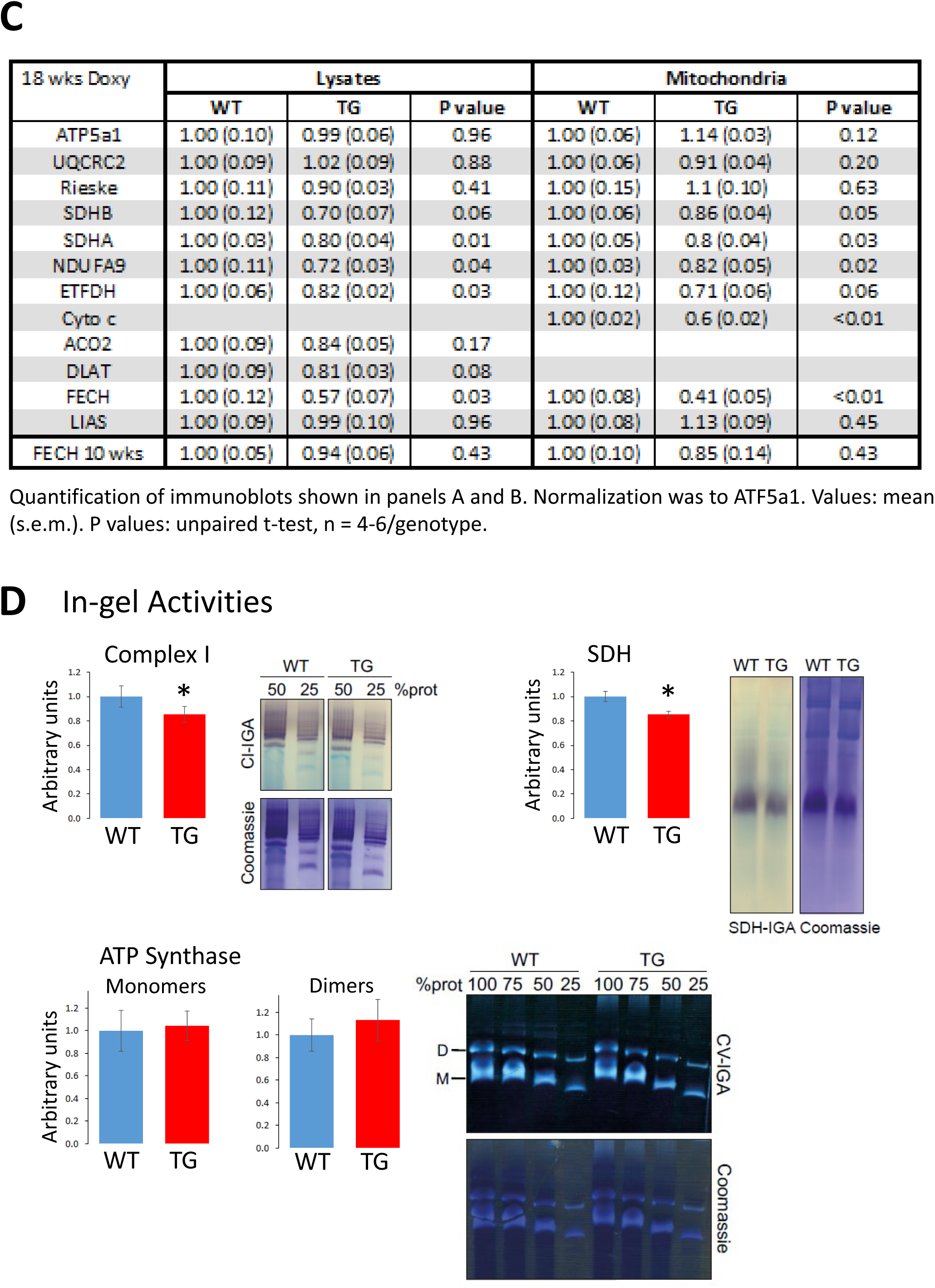
Selected immunoblot analysis of mitochondrial proteins. **(A)** Examples of immunoblots from heart lysates and mitochondria from mice fed Doxy for 18 wks **(B)** Examples of immunoblots from heart mitochondria from mice fed Doxy for 10 wks. In A and B, bold lettering refers to proteins that contain ISCs, heme or lipoic acid. *: proteins showing significant changes in TG samples (see quantification in **C**) **(C)** Quantification of immunoblots from heart lysates and mitochondria from 18-wk Doxy fed mice. Values were normalized to ATP5a1. Values are the mean ± s.e.m. P values: unpaired t-test, n=4/genotype for lysates, n=5/genotype for mitochondria. **(D)** In-gel activity for Complex I, SDH and ATP Synthase. Values are the mean ± s.e.m. * p=0.03, paired t-test; Complex I and SDH: n=5/genotype; ATP Synthase: n=3/genotype.

**Figure S4:**
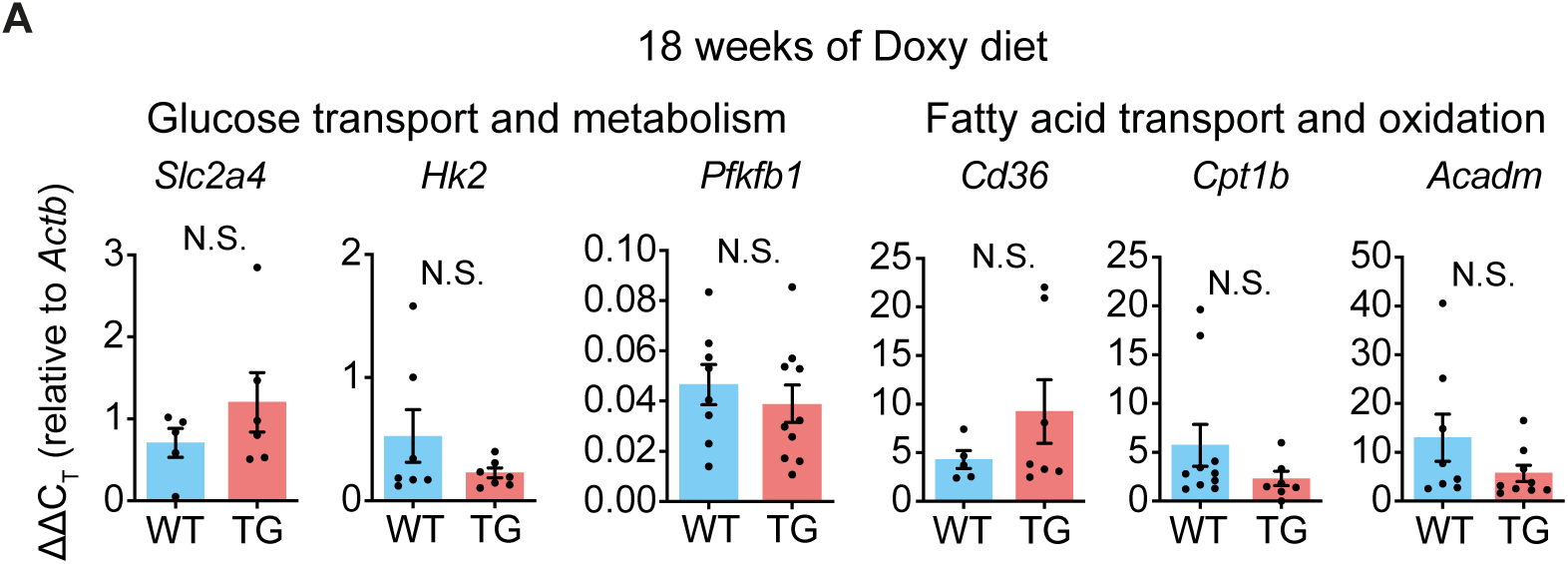
No change in transcript levels of genes encoding glycolytic and fatty acid handling and metabolism proteins in hearts from TG mice fed Doxy diet for 18 weeks. ΔΔCT values of transcripts involved in fatty acid and glucose oxidation and transport, normalized to *Actb* (β-actin). Bars represent the mean ± s.e.m; individual points represent values from each mouse. Statistical analysis was by unpaired t-test, N.S.: not significant, n=8/genotype.

**Figure S5:**
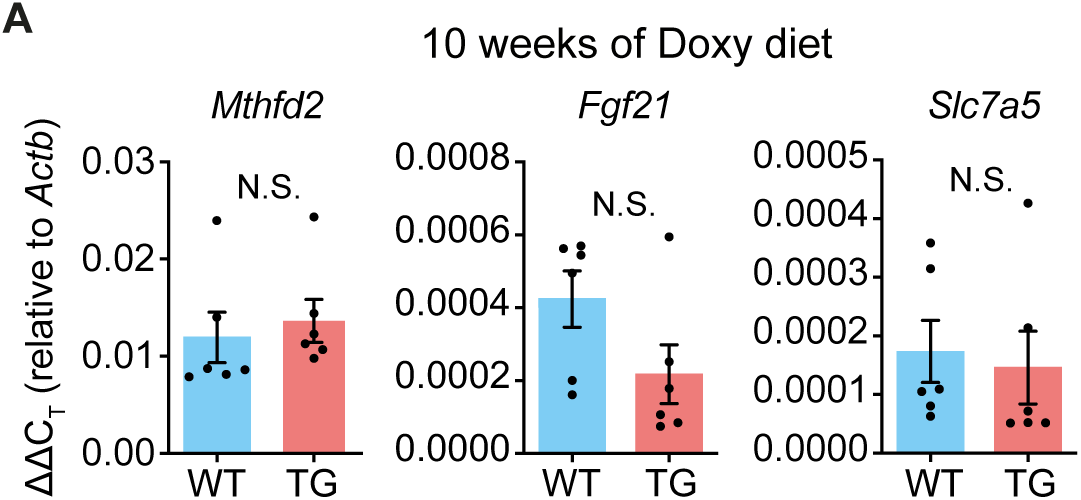
No change in ATF4 targets in the heart after 10 wks on Doxy diet. *Actb* (β-Actin) normalized transcripts (expressed as ΔΔCT) of ATF4 target genes in mice fed Doxy diet for 10 weeks. Bars represent the mean ± s.e.m; individual points represent values from each mouse. Statistical analysis: unpaired t-test, N.S.: not significant, n=4/genotype.

**Figure S6:**
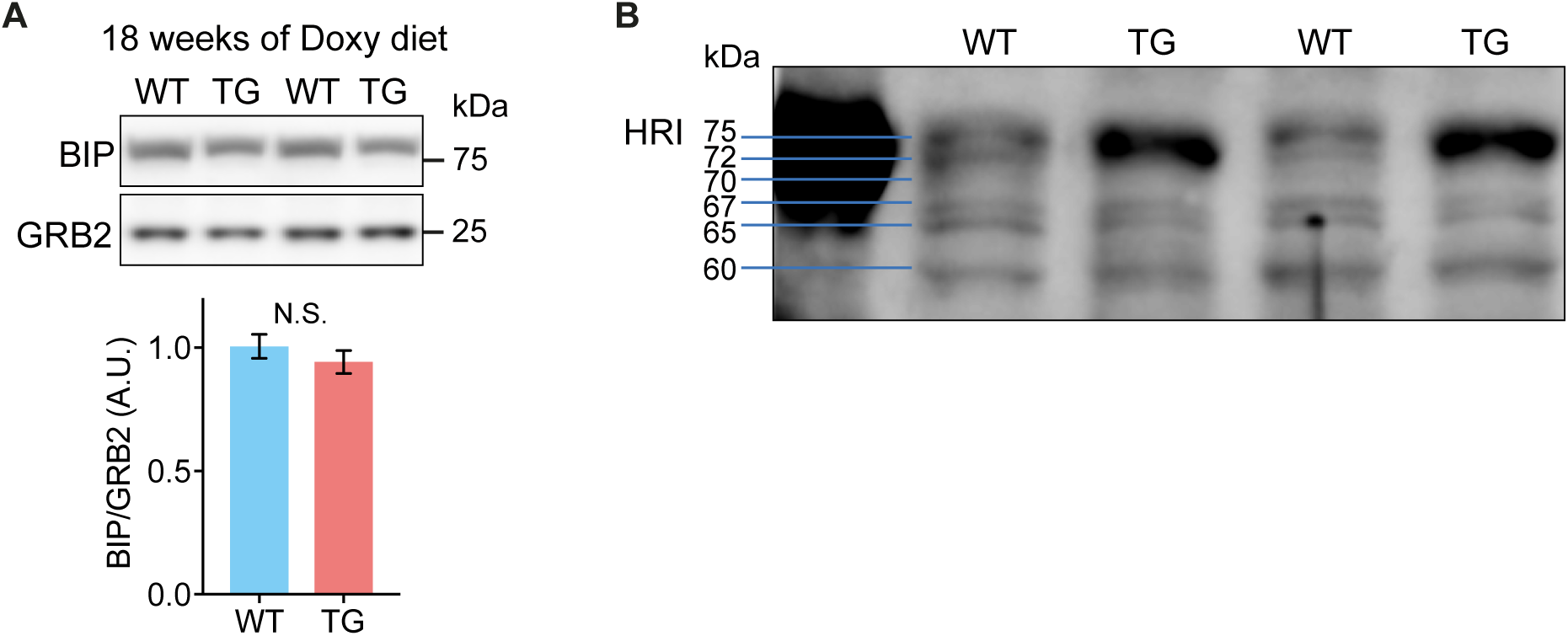
Absence of endoplasmic reticulum stress in the heart after 18 wks of Doxy diet. **(A)** *Upper*: Representative immunoblot of endoplasmic reticulum chaperon protein BIP, and GRB2 (loading control) from heart lysates of mice fed Doxy diet for 18 wks, and quantification (*lower*), means +/- sem. Statistical comparison: unpaired t-test, N.S.: not significant. (n=9 WT/7 TG). **(B)** Calibration of immunoblot using anti-HRI antibody.

## BIBLIOGRAPHY

1. Reetz, K., Dogan, I., Costa, A.S., Dafotakis, M., Fedosov, K., Giunti, P., Parkinson, M.H., Sweeney, M.G., Mariotti, C., Panzeri, M. et al. (2015) Biological and clinical characteristics of the European Friedreich’s Ataxia Consortium for Translational Studies (EFACTS) cohort: a cross-sectional analysis of baseline data. Lancet Neurol, 14, 174–182.

2. Kipps, A., Alexander, M., Colan, S.D., Gauvreau, K., Smoot, L., Crawford, L., Darras, B.T. and Blume, E.D. (2009) The longitudinal course of cardiomyopathy in Friedreich’s ataxia during childhood. Pediatr Cardiol, 30, 306–310.

3. Tsou, A.Y., Paulsen, E.K., Lagedrost, S.J., Perlman, S.L., Mathews, K.D., Wilmot, G.R., Ravina, B., Koeppen, A.H. and Lynch, D.R. (2011) Mortality in Friedreich ataxia. J Neurol Sci, 307, 46–49.

4. St John Sutton, M., Ky, B., Regner, S.R., Schadt, K., Plappert, T., He, J., D’Souza, B. and Lynch, D.R. (2014) Longitudinal strain in Friedreich Ataxia: a potential marker for early left ventricular dysfunction. Echocardiography, 31, 50–57.

5. Legrand, L., Diallo, A., Monin, M.L., Ewenczyk, C., Charles, P., Isnard, R., Vicaut, E., Montalescot, G., Durr, A. and Pousset, F. (2020) Predictors of Left Ventricular Dysfunction in Friedreich’s Ataxia in a 16-Year Observational Study. Am J Cardiovasc Drugs, 20, 209–216.

6. Huang, M.L., Sivagurunathan, S., Ting, S., Jansson, P.J., Austin, C.J., Kelly, M., Semsarian, C., Zhang, D. and Richardson, D.R. (2013) Molecular and functional alterations in a mouse cardiac model of Friedreich ataxia: activation of the integrated stress response, eIF2alpha phosphorylation, and the induction of downstream targets. Am J Pathol, 183, 745–757.

7. Pakos-Zebrucka, K., Koryga, I., Mnich, K., Ljujic, M., Samali, A. and Gorman, A.M. (2016) The integrated stress response. EMBO Rep, 17, 1374–1395.

8. Costa-Mattioli, M. and Walter, P. (2020) The integrated stress response: From mechanism to disease. Science, 368.

9. Suomalainen, A., Elo, J.M., Pietilainen, K.H., Hakonen, A.H., Sevastianova, K., Korpela, M., Isohanni, P., Marjavaara, S.K., Tyni, T., Kiuru-Enari, S. et al. (2011) FGF-21 as a biomarker for muscle-manifesting mitochondrial respiratory chain deficiencies: a diagnostic study. Lancet Neurol, 10, 806–818.

10. Bao, X.R., Ong, S.E., Goldberger, O., Peng, J., Sharma, R., Thompson, D.A., Vafai, S.B., Cox, A.G., Marutani, E., Ichinose, F. et al. (2016) Mitochondrial dysfunction remodels one-carbon metabolism in human cells. Elife, 5.

11. Nikkanen, J., Forsstrom, S., Euro, L., Paetau, I., Kohnz, R.A., Wang, L., Chilov, D., Viinamaki, J., Roivainen, A., Marjamaki, P. et al. (2016) Mitochondrial DNA Replication Defects Disturb Cellular dNTP Pools and Remodel One-Carbon Metabolism. Cell Metab, 23, 635–648.

12. Khan, N.A., Nikkanen, J., Yatsuga, S., Jackson, C., Wang, L., Pradhan, S., Kivela, R., Pessia, A., Velagapudi, V. and Suomalainen, A. (2017) mTORC1 Regulates Mitochondrial Integrated Stress Response and Mitochondrial Myopathy Progression. Cell Metab, 26, 419–428 e415.

13. Quiros, P.M., Prado, M.A., Zamboni, N., D’Amico, D., Williams, R.W., Finley, D., Gygi, S.P. and Auwerx, J. (2017) Multi-omics analysis identifies ATF4 as a key regulator of the mitochondrial stress response in mammals. J Cell Biol, 216, 2027–2045.

14. Dogan, S.A., Cerutti, R., Beninca, C., Brea-Calvo, G., Jacobs, H.T., Zeviani, M., Szibor, M. and Viscomi, C. (2018) Perturbed Redox Signaling Exacerbates a Mitochondrial Myopathy. Cell Metab, 28, 764–775 e765.

15. Forsstrom, S., Jackson, C.B., Carroll, C.J., Kuronen, M., Pirinen, E., Pradhan, S., Marmyleva, A., Auranen, M., Kleine, I.M., Khan, N.A. et al. (2019) Fibroblast Growth Factor 21 Drives Dynamics of Local and Systemic Stress Responses in Mitochondrial Myopathy with mtDNA Deletions. Cell Metab, 30, 1040–1054 e1047.

16. Mick, E., Titov, D.V., Skinner, O.S., Sharma, R., Jourdain, A.A. and Mootha, V.K. (2020) Distinct mitochondrial defects trigger the integrated stress response depending on the metabolic state of the cell. Elife, 9.

17. Johnson, S.C., Yanos, M.E., Kayser, E.B., Quintana, A., Sangesland, M., Castanza, A., Uhde, L., Hui, J., Wall, V.Z., Gagnidze, A. et al. (2013) mTOR inhibition alleviates mitochondrial disease in a mouse model of Leigh syndrome. Science, 342, 1524–1528.

18. Siegmund, S.E., Yang, H., Sharma, R., Javors, M., Skinner, O., Mootha, V., Hirano, M. and Schon, E.A. (2017) Low-dose rapamycin extends lifespan in a mouse model of mtDNA depletion syndrome. Hum Mol Genet, 26, 4588–4605.

19. Civiletto, G., Dogan, S.A., Cerutti, R., Fagiolari, G., Moggio, M., Lamperti, C., Beninca, C., Viscomi, C. and Zeviani, M. (2018) Rapamycin rescues mitochondrial myopathy via coordinated activation of autophagy and lysosomal biogenesis. EMBO Mol Med, 10.

20. Viscomi, C., Bottani, E., Civiletto, G., Cerutti, R., Moggio, M., Fagiolari, G., Schon, E.A., Lamperti, C. and Zeviani, M. (2011) In vivo correction of COX deficiency by activation of the AMPK/PGC-1alpha axis. Cell Metab, 14, 80–90.

21. Seznec, H., Simon, D., Monassier, L., Criqui-Filipe, P., Gansmuller, A., Rustin, P., Koenig, M. and Puccio, H. (2004) Idebenone delays the onset of cardiac functional alteration without correction of Fe-S enzymes deficit in a mouse model for Friedreich ataxia. Hum Mol Genet, 13, 1017–1024.

22. Wagner, G.R., Pride, P.M., Babbey, C.M. and Payne, R.M. (2012) Friedreich’s ataxia reveals a mechanism for coordinate regulation of oxidative metabolism via feedback inhibition of the SIRT3 deacetylase. Hum Mol Genet, 21, 2688–2697.

23. Stram, A.R., Wagner, G.R., Fogler, B.D., Pride, P.M., Hirschey, M.D. and Payne, R.M. (2017) Progressive mitochondrial protein lysine acetylation and heart failure in a model of Friedreich’s ataxia cardiomyopathy. PLoS One, 12, e0178354.

24. 24 Martin, A.S., Abraham, D.M., Hershberger, K.A., Bhatt, D.P., Mao, L., Cui, H., Liu, J., Liu, X., Muehlbauer, M.J., Grimsrud, P.A. et al. (2017) Nicotinamide mononucleotide requires SIRT3 to improve cardiac function and bioenergetics in a Friedreich’s ataxia cardiomyopathy model. JCI Insight, 2.

25. Perdomini, M., Belbellaa, B., Monassier, L., Reutenauer, L., Messaddeq, N., Cartier, N., Crystal, R.G., Aubourg, P. and Puccio, H. (2014) Prevention and reversal of severe mitochondrial cardiomyopathy by gene therapy in a mouse model of Friedreich’s ataxia. Nat Med, 20, 542–547.

26. Belbellaa, B., Reutenauer, L., Monassier, L. and Puccio, H. (2019) Correction of half the cardiomyocytes fully rescue Friedreich ataxia mitochondrial cardiomyopathy through cell-autonomous mechanisms. Hum Mol Genet, 28, 1274–1285.

27. Chandran, V., Gao, K., Swarup, V., Versano, R., Dong, H., Jordan, M.C. and Geschwind, D.H. (2017) Inducible and reversible phenotypes in a novel mouse model of Friedreich’s Ataxia. Elife, 6.

28. Llorens, J.V., Soriano, S., Calap-Quintana, P., Gonzalez-Cabo, P. and Molto, M.D. (2019) The Role of Iron in Friedreich’s Ataxia: Insights From Studies in Human Tissues and Cellular and Animal Models. Front Neurosci, 13, 75.

29. Anderson, C.P., Shen, M., Eisenstein, R.S. and Leibold, E.A. (2012) Mammalian iron metabolism and its control by iron regulatory proteins. Biochim Biophys Acta, 1823, 1468–1483.

30. Razeghi, P., Young, M.E., Alcorn, J.L., Moravec, C.S., Frazier, O.H. and Taegtmeyer, H. (2001) Metabolic gene expression in fetal and failing human heart. Circulation, 104, 2923–2931.

31. Wojtczak, L. and Schonfeld, P. (1993) Effect of fatty acids on energy coupling processes in mitochondria. Biochim Biophys Acta, 1183, 41–57.

32. Branda, S.S., Cavadini, P., Adamec, J., Kalousek, F., Taroni, F. and Isaya, G. (1999) Yeast and human frataxin are processed to mature form in two sequential steps by the mitochondrial processing peptidase. J Biol Chem, 274, 22763–22769.

33. Bertholet, A.M., Chouchani, E.T., Kazak, L., Angelin, A., Fedorenko, A., Long, J.Z., Vidoni, S., Garrity, R., Cho, J., Terada, N. et al. (2019) H(+) transport is an integral function of the mitochondrial ADP/ATP carrier. Nature, 571, 515–520.

34. Vinothkumar, K.R., Zhu, J. and Hirst, J. (2014) Architecture of mammalian respiratory complex I. Nature, 515, 80–84.

35. Zhu, J., Vinothkumar, K.R. and Hirst, J. (2016) Structure of mammalian respiratory complex I. Nature, 536, 354–358.

36. Ndi, M., Marin-Buera, L., Salvatori, R., Singh, A.P. and Ott, M. (2018) Biogenesis of the bc1 Complex of the Mitochondrial Respiratory Chain. J Mol Biol, 430, 3892–3905.

37. Wende, A.R., Brahma, M.K., McGinnis, G.R. and Young, M.E. (2017) Metabolic Origins of Heart Failure. JACC Basic Transl Sci, 2, 297–310.

38. Chung, H.K., Ryu, D., Kim, K.S., Chang, J.Y., Kim, Y.K., Yi, H.S., Kang, S.G., Choi, M.J., Lee, S.E., Jung, S.B. et al. (2017) Growth differentiation factor 15 is a myomitokine governing systemic energy homeostasis. J Cell Biol, 216, 149–165.

39. Kuhl, I., Miranda, M., Atanassov, I., Kuznetsova, I., Hinze, Y., Mourier, A., Filipovska, A. and Larsson, N.G. (2017) Transcriptomic and proteomic landscape of mitochondrial dysfunction reveals secondary coenzyme Q deficiency in mammals. Elife, 6.

40. Park, Y., Reyna-Neyra, A., Philippe, L. and Thoreen, C.C. (2017) mTORC1 Balances Cellular Amino Acid Supply with Demand for Protein Synthesis through Post-transcriptional Control of ATF4. Cell Rep, 19, 1083–1090.

41. Selvarajah, B., Azuelos, I., Plate, M., Guillotin, D., Forty, E.J., Contento, G., Woodcock, H.V., Redding, M., Taylor, A., Brunori, G. et al. (2019) mTORC1 amplifies the ATF4-dependent de novo serine-glycine pathway to supply glycine during TGF-beta1-induced collagen biosynthesis. Sci Signal, 12.

42. Kovacic, S., Soltys, C.L., Barr, A.J., Shiojima, I., Walsh, K. and Dyck, J.R. (2003) Akt activity negatively regulates phosphorylation of AMP-activated protein kinase in the heart. J Biol Chem, 278, 39422–39427.

43. Soltys, C.L., Kovacic, S. and Dyck, J.R. (2006) Activation of cardiac AMP-activated protein kinase by LKB1 expression or chemical hypoxia is blunted by increased Akt activity. Am J Physiol Heart Circ Physiol, 290, H2472–2479.

44. Suragani, R.N., Zachariah, R.S., Velazquez, J.G., Liu, S., Sun, C.W., Townes, T.M. and Chen, J.J. (2012) Heme-regulated eIF2alpha kinase activated Atf4 signaling pathway in oxidative stress and erythropoiesis. Blood, 119, 5276–5284.

45. Zhang, S., Macias-Garcia, A., Velazquez, J., Paltrinieri, E., Kaufman, R.J. and Chen, J.J. (2018) HRI coordinates translation by eIF2alphaP and mTORC1 to mitigate ineffective erythropoiesis in mice during iron deficiency. Blood, 131, 450–461.

46. Schmidt, E.K., Clavarino, G., Ceppi, M. and Pierre, P. (2009) SUnSET, a nonradioactive method to monitor protein synthesis. Nat Methods, 6, 275–277.

47. Kanamori, H., Takemura, G., Maruyama, R., Goto, K., Tsujimoto, A., Ogino, A., Li, L., Kawamura, I., Takeyama, T., Kawaguchi, T. et al. (2009) Functional significance and morphological characterization of starvation-induced autophagy in the adult heart. Am J Pathol, 174, 1705–1714.

48. Andres, A.M., Kooren, J.A., Parker, S.J., Tucker, K.C., Ravindran, N., Ito, B.R., Huang, C., Venkatraman, V., Van Eyk, J.E., Gottlieb, R.A. et al. (2016) Discordant signaling and autophagy response to fasting in hearts of obese mice: Implications for ischemia tolerance. Am J Physiol Heart Circ Physiol, 311, H219–228.

49. Gubbiotti, M.A., Seifert, E., Rodeck, U., Hoek, J.B. and Iozzo, R.V. (2018) Metabolic reprogramming of murine cardiomyocytes during autophagy requires the extracellular nutrient sensor decorin. J Biol Chem, 293, 16940–16950.

50. Willis, M.S., Rojas, M., Li, L., Selzman, C.H., Tang, R.H., Stansfield, W.E., Rodriguez, J.E., Glass, D.J. and Patterson, C. (2009) Muscle ring finger 1 mediates cardiac atrophy in vivo. Am J Physiol Heart Circ Physiol, 296, H997–H1006.

51. Willis, M.S., Parry, T.L., Brown, D.I., Mota, R.I., Huang, W., Beak, J.Y., Sola, M., Zhou, C., Hicks, S.T., Caughey, M.C. et al. (2019) Doxorubicin Exposure Causes Subacute Cardiac Atrophy Dependent on the Striated Muscle-Specific Ubiquitin Ligase MuRF1. Circ Heart Fail, 12, e005234.

52. Mochel, F., Knight, M.A., Tong, W.H., Hernandez, D., Ayyad, K., Taivassalo, T., Andersen, P.M., Singleton, A., Rouault, T.A., Fischbeck, K.H. et al. (2008) Splice mutation in the iron-sulfur cluster scaffold protein ISCU causes myopathy with exercise intolerance. Am J Hum Genet, 82, 652–660.

53. Rotig, A., de Lonlay, P., Chretien, D., Foury, F., Koenig, M., Sidi, D., Munnich, A. and Rustin, P. (1997) Aconitase and mitochondrial iron-sulphur protein deficiency in Friedreich ataxia. Nat Genet, 17, 215–217.

54. Navarro-Sastre, A., Tort, F., Stehling, O., Uzarska, M.A., Arranz, J.A., Del Toro, M., Labayru, M.T., Landa, J., Font, A., Garcia-Villoria, J. et al. (2011) A fatal mitochondrial disease is associated with defective NFU1 function in the maturation of a subset of mitochondrial Fe-S proteins. Am J Hum Genet, 89, 656–667.

55. Lim, S.C., Friemel, M., Marum, J.E., Tucker, E.J., Bruno, D.L., Riley, L.G., Christodoulou, J., Kirk, E.P., Boneh, A., DeGennaro, C.M. et al. (2013) Mutations in LYRM4, encoding iron-sulfur cluster biogenesis factor ISD11, cause deficiency of multiple respiratory chain complexes. Hum Mol Genet, 22, 4460–4473.

56. Crooks, D.R., Maio, N., Lane, A.N., Jarnik, M., Higashi, R.M., Haller, R.G., Yang, Y., Fan, T.W., Linehan, W.M. and Rouault, T.A. (2018) Acute loss of iron-sulfur clusters results in metabolic reprogramming and generation of lipid droplets in mammalian cells. J Biol Chem, 293, 8297–8311.

57. Fornasiero, E.F., Mandad, S., Wildhagen, H., Alevra, M., Rammner, B., Keihani, S., Opazo, F., Urban, I., Ischebeck, T., Sakib, M.S. et al. (2018) Precisely measured protein lifetimes in the mouse brain reveal differences across tissues and subcellular fractions. Nat Commun, 9, 4230.

58. Szczepanowska, K., Senft, K., Heidler, J., Herholz, M., Kukat, A., Hohne, M.N., Hofsetz, E., Becker, C., Kaspar, S., Giese, H. et al. (2020) A salvage pathway maintains highly functional respiratory complex I. Nat Commun, 11, 1643.

59. Mik, E.G., Ince, C., Eerbeek, O., Heinen, A., Stap, J., Hooibrink, B., Schumacher, C.A., Balestra, G.M., Johannes, T., Beek, J.F. et al. (2009) Mitochondrial oxygen tension within the heart. J Mol Cell Cardiol, 46, 943–951.

60. Ast, T., Meisel, J.D., Patra, S., Wang, H., Grange, R.M.H., Kim, S.H., Calvo, S.E., Orefice, L.L., Nagashima, F., Ichinose, F. et al. (2019) Hypoxia Rescues Frataxin Loss by Restoring Iron Sulfur Cluster Biogenesis. Cell, 177, 1507–1521 e1516.

61. Tyynismaa, H., Carroll, C.J., Raimundo, N., Ahola-Erkkila, S., Wenz, T., Ruhanen, H., Guse, K., Hemminki, A., Peltola-Mjosund, K.E., Tulkki, V. et al. (2010) Mitochondrial myopathy induces a starvation-like response. Hum Mol Genet, 19, 3948–3958.

62. Liu, G.Y. and Sabatini, D.M. (2020) mTOR at the nexus of nutrition, growth, ageing and disease. Nat Rev Mol Cell Biol, 21, 183–203.

63. Varghese, J., James, J., Vaulont, S., McKie, A. and Jacob, M. (2018) Increased intracellular iron in mouse primary hepatocytes in vitro causes activation of the Akt pathway but decreases its response to insulin. Biochim Biophys Acta Gen Subj, 1862, 1870–1882.

64. Ellis, J.M., Mentock, S.M., Depetrillo, M.A., Koves, T.R., Sen, S., Watkins, S.M., Muoio, D.M., Cline, G.W., Taegtmeyer, H., Shulman, G.I. et al. (2011) Mouse cardiac acyl coenzyme a synthetase 1 deficiency impairs Fatty Acid oxidation and induces cardiac hypertrophy. Mol Cell Biol, 31, 1252–1262.

65. Hawley, S.A., Ross, F.A., Gowans, G.J., Tibarewal, P., Leslie, N.R. and Hardie, D.G. (2014) Phosphorylation by Akt within the ST loop of AMPK-alpha1 down-regulates its activation in tumour cells. Biochem J, 459, 275–287.

66. Han, A.P., Yu, C., Lu, L., Fujiwara, Y., Browne, C., Chin, G., Fleming, M., Leboulch, P., Orkin, S.H. and Chen, J.J. (2001) Heme-regulated eIF2alpha kinase (HRI) is required for translational regulation and survival of erythroid precursors in iron deficiency. EMBO J, 20, 6909–6918.

67. Guo, X., Aviles, G., Liu, Y., Tian, R., Unger, B.A., Lin, Y.T., Wiita, A.P., Xu, K., Correia, M.A. and Kampmann, M. (2020) Mitochondrial stress is relayed to the cytosol by an OMA1-DELE1-HRI pathway. Nature, 579, 427–432.

68. Fessler, E., Eckl, E.M., Schmitt, S., Mancilla, I.A., Meyer-Bender, M.F., Hanf, M., Philippou-Massier, J., Krebs, S., Zischka, H. and Jae, L.T. (2020) A pathway coordinated by DELE1 relays mitochondrial stress to the cytosol. Nature, 579, 433–437.

69. Pereira, R.O., Tadinada, S.M., Zasadny, F.M., Oliveira, K.J., Pires, K.M.P., Olvera, A., Jeffers, J., Souvenir, R., McGlauflin, R., Seei, A. et al. (2017) OPA1 deficiency promotes secretion of FGF21 from muscle that prevents obesity and insulin resistance. EMBO J, 36, 2126–2145.

70. Moffat, C., Bhatia, L., Nguyen, T., Lynch, P., Wang, M., Wang, D., Ilkayeva, O.R., Han, X., Hirschey, M.D., Claypool, S.M. et al. (2014) Acyl-CoA thioesterase-2 facilitates mitochondrial fatty acid oxidation in the liver. J Lipid Res, 55, 2458–2470.

71. Jha, P., Wang, X. and Auwerx, J. (2016) Analysis of Mitochondrial Respiratory Chain Supercomplexes Using Blue Native Polyacrylamide Gel Electrophoresis (BN-PAGE). Curr Protoc Mouse Biol, 6, 1–14.

72. Cox, J. and Mann, M. (2008) MaxQuant enables high peptide identification rates, individualized p.p.b.-range mass accuracies and proteome-wide protein quantification. Nat Biotechnol, 26, 1367–1372.

73. Tyanova, S., Temu, T., Sinitcyn, P., Carlson, A., Hein, M.Y., Geiger, T., Mann, M. and Cox, J. (2016) The Perseus computational platform for comprehensive analysis of (prote)omics data. Nat Methods, 13, 731–740.

